# Leveraging Large Language Models for Redundancy-Aware Pathway Analysis and Deep Biological Interpretation

**DOI:** 10.1101/2025.08.23.671949

**Authors:** Yifei Ge, Feifan Zhang, Yijiang Liu, Chao Jiang, Peng Gao, Nguan Soon Tan, Sai Zhang, Yuchen Shen, Qianyi Zhou, Xin Zhou, Chuchu Wang, Xiaotao Shen

## Abstract

Extracting coherent, biologically meaningful insights from vast, complex multi-omics data remains challenging. Currently, pathway enrichment analysis serves as a cornerstone for the functional interpretation of such data. However, conventional approaches often suffer from extensive functional redundancy caused by shared molecular components and overlapping pathway definitions across databases. This redundancy can obscure key biological signals and compromise the interpretability of pathway enrichment results. Here, we present MAPA (Functional **M**odule Identification and **A**nnotation for **P**athway **A**nalysis Results Using Large Language Models [LLM]), an open-source computational framework that resolves redundancy and enhances pathway analysis result interpretation. MAPA computes functional similarity between pathways using LLM-based text embeddings, enabling comparison across different databases. It constructs pathway similarity networks and identifies functional modules via community detection algorithms. Crucially, MAPA employs LLMs for automated functional annotation, integrating Retrieval-Augmented Generation (RAG) to generate comprehensive and real-time biological summaries and reduce hallucinations. Benchmarking demonstrated MAPA’s superior performance: the biotext embedding similarity showed a large effect size (Cliff’s δ = 0.96) compared with the Jaccard index (δ = 0.73), and module identification achieved high accuracy (Adjusted Rand Index [ARI] = 0.95) versus existing methods (ARI = 0.23-0.33). Human expert evaluation confirmed that MAPA’s annotations match expert-quality interpretations. Finally, a multi-omics aging case study illustrates that MAPA uncovers coherent functional modules and generates insights extending beyond conventional pathway analyses. Collectively, MAPA represents a significant advance in redundancy-aware pathway analysis, transforming pathway enrichment results from fragmented lists into biologically coherent narratives. By leveraging the capabilities of LLMs, MAPA offers researchers a robust, scalable tool for deriving deep mechanistic insights from complex and vast multi-omics datasets, marking a new direction for AI-driven bioinformatics.

## Introduction

Multi-omics technologies have become indispensable tools in fundamental biomedical research^1–6^. A central challenge in omics research is to identify groups of biomarkers and translate them into biologically meaningful insights. Currently, one of the most widely used strategies is pathway analysis, which aims to contextualize biomarkers within the framework of known biological processes^7–9^. By leveraging these approaches, researchers can infer the biological functions associated with biomarkers, enabling a deeper and more comprehensive understanding of the molecular mechanisms underpinning various physiological states and disease conditions^3,10–13^. However, these analyses often yield results burdened by significant functional redundancy (pathways that have the same or similar biological functions), driven by overlapping pathway definitions within and across databases^3,11,12^. Such redundancy complicates interpretation, frequently obscuring biologically meaningful insights when results are reported simply as the top N enriched pathways^14–17^. Addressing redundancy is thus crucial for enhancing the clarity, biological relevance, and actionable value of multi-omics data.

To address the pervasive issue of redundancy in pathway analysis results, a range of computational approaches has been developed to improve the interpretability and biological relevance of high-dimensional omics data^18–35^. For instance, REVIGO clusters Gene Ontology (GO) terms based on semantic similarity, allowing users to retain a subset of representative, non-redundant terms^19^. Similarly, Enrichment Map, a Cytoscape plugin, constructs similarity networks among enriched GO terms to visualize functionally coherent modules and reveal broader biological themes^20^. Other tools, such as enrichplot^36^ and aPEAR^26^, offer solutions that filter highly similar pathways by computing quantitative similarity metrics like the Jaccard index^37^. Additionally, methods like topGO incorporate graph-based algorithms to account for hierarchical relationships within the GO structure^38^. These methods have become integral to current omics workflows, providing critical capabilities for reducing redundancy and summarizing complex enrichment results into more interpretable biological narratives^3,15,39–41^.

However, existing redundancy-reduction strategies exhibit several key limitations. First, many approaches depend heavily on semantic similarity metrics specifically optimized for GO terms^7,19,23,30,31^, limiting their applicability to other pathway databases such as Kyoto Encyclopedia of Genes and Genomes (KEGG)^42^, Reactome^43^, WikiPathways^44^, and Small Molecule Pathway Database (SMPDB)^45^. Second, most tools do not provide a systematic functional interpretation of the identified modules, thereby failing to capture the collective biological context within each module^18,26–28^. Third, redundancy reduction is often treated as a post hoc visualization or filtering step rather than being integrated into the core systemic framework of pathway analysis, which restricts scalability and interpretability.

Recent advances in LLMs present a unique opportunity to address these limitations. Trained on vast biomedical corpora, LLMs can capture the nuanced semantic connections (functional similarity) between pathways that lie beyond rigid ontologies or simple gene-overlap counts^46^. This capability, combined with their ability to generate coherent summaries and integrate diverse information sources^47–49^, makes them particularly well-suited for providing interpretable functional annotations. Furthermore, recent advances in retrieval-augmented generation (RAG) further strengthen LLMs by allowing models to query the latest literature on demand, keeping annotations current while curbing hallucinations common to purely generative systems^1,50,51^. In the biomedical domain, LLMs have been successfully applied to diverse tasks, including biomedical literature summarization^46^, gene set annotation^47,49^, and extraction of relationships between biomedical data and diseases^48^. Together with these capabilities, LLM-based workflows offer unprecedented opportunities for enhancing data interpretation in the multi-omics field.

Here, we introduce MAPA (Functional **M**odule Identification and **A**nnotation for **P**athway **A**nalysis Results Using LLMs), a novel computational framework designed to systematically reduce pathway redundancy and enhance the interpretability of multi-omics data. MAPA computes functional similarities among enriched pathways using LLM-based biological text embeddings. This allows MAPA to capture nuanced functional similarities of pathways across different databases. These biological text embedding similarities are used to construct a pathway similarity network, from which MAPA identifies functional modules. Crucially, MAPA employs LLMs for automated and unbiased functional annotation. By leveraging RAG, MAPA utilizes relevant, real-time literature to reduce hallucinations and generate accurate and comprehensive biological summaries for each module. Finally, we demonstrate the power of MAPA in a multi-omics case study of aging, showing how it uncovers coherent functional modules and delivers rich biological interpretations that would remain hidden using conventional pathway analysis approaches. Together, MAPA represents a significant advance in redundancy-aware pathway analysis, providing a scalable and interpretable solution for extracting biologically meaningful insights from vast and complex multi-omics data.

## Results

### Pathway Functional Redundancy Is Pervasive Within and Across Pathway Databases

Various studies have shown that pathway functional redundancy is pervasive within and across pathway databases^29,34,52^. To systematically assess the extent of pathway redundancy that can obscure biological interpretation, we first analyzed the internal structure of individual pathway databases (**Methods**). Focusing on the GO Biological Process (BP) ontology, we calculated pairwise semantic similarity among all GO terms using the Wang method^53^. The resulting distribution revealed widespread redundancy, with a substantial fraction of term pairs exhibiting high semantic similarity (**Fig. 1a**). Similar patterns were observed for the Molecular Function (MF) and Cellular Component (CC) branches of GO (**Supplementary Figure 1a,b**), demonstrating that redundancy is a pervasive feature within pathway databases.

**Figure 1.**
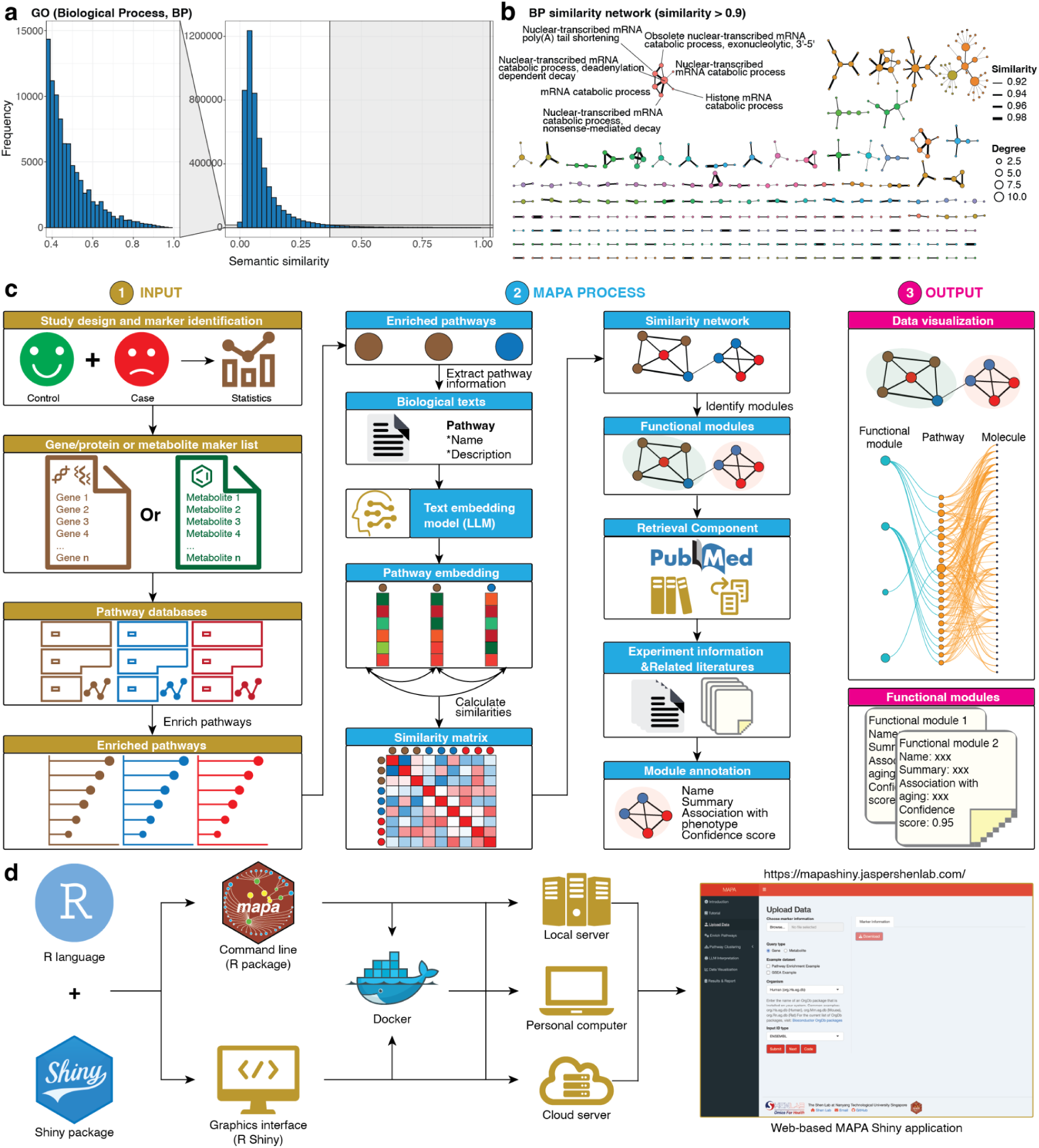
Redundancy in pathway databases and the design of the MAPA framework. (**a**) Distribution of pairwise semantic similarities among GO BP terms. A substantial proportion of GO term pairs exhibit high semantic similarity, indicating significant within-database redundancy. (**b**) The similarity network of GO BP terms, constructed using a similarity threshold of > 0.9. Each node represents a GO term, and edges indicate high semantic similarity. A representative module is highlighted, consisting of tightly clustered terms related to nuclear-transcribed mRNA decay. (**c**) Overview of the MAPA framework. MAPA takes enriched pathways as input, calculates multi-modal similarity based on biological text embeddings, builds a similarity network, identifies functional modules, and uses LLMs to generate human-interpretable annotations for each functional module. (**d**) Development and deployment of the MAPA framework, including integration with the R ecosystem, web-based interface, and availability as an open-source tool with support for local or server-based applications.

To further visualize the structure of this redundancy, we constructed a similarity network for GO BP terms using a threshold of semantic similarity > 0.9 (**Methods**). This network resolved into numerous tightly connected modules (**Fig. 1b**), each comprising terms with highly overlapping biological meaning (**Supplementary Data 1**). For example, one module included the GO terms Nuclear-transcribed mRNA poly(A) tail shortening, Nuclear-transcribed mRNA catabolic process, deadenylation-dependent decay, nonsense-mediated decay, and related processes (**Fig. 1b**). This module collectively describes the nuclear-transcribed mRNA decay machinery, which governs transcript turnover via deadenylation and exonucleolytic degradation pathways. The clear functional coherence of this module, as well as its dense intra-module similarity and lack of inter-module connections, highlights the redundancy among functionally similar GO terms within the database (**Fig. 1b**).

We next extended this analysis to examine cross-database redundancy. By computing the pairwise Jaccard index based on molecule content between pathways from three different metabolic databases (**Methods**), we observed extensive overlap across databases (**Supplementary Figure 1c-e**). Many biological processes, such as glycolysis and fatty acid metabolism, were represented by overlapping pathways across databases, each annotated differently in detailed descriptions but sharing a large proportion of molecular components (**Supplementary Figure 1f**).

Together, all these results demonstrate that both within- and across-database redundancy are widespread and intrinsic to existing popular pathway databases. This redundancy complicates the interpretation of pathways, underscoring the need for a systematic framework to resolve, cluster, and annotate overlapping pathways into coherent functional modules.

### MAPA Workflow

To address the pervasive functional redundancy within and across pathway databases, we developed MAPA, a computational framework designed to transform results of pathway enrichment analysis into coherent and interpretable functional modules (**Fig. 1c**). MAPA is compatible with standard pathway enrichment methods, including Overrepresentation Analysis (ORA)^54^ and Gene Set Enrichment Analysis (GSEA)^8^, and supports multi-omics inputs from transcriptomics, proteomics, and metabolomics studies (**Methods**). Users provide a list of differentially expressed or ranked genes, proteins, or metabolites, which are subjected to enrichment analysis in MAPA using pathway databases (**Fig. 1c**).

The enriched pathways are then passed to the functional module identification pipeline (**Fig. 1c** and **Methods**). For each pathway, MAPA extracts relevant biological text, specifically the pathway name and functional description, and encodes this information using an LLM-based text embedding model. Each pathway is thereby converted into a high-dimensional vector that captures its functional and semantic characteristics. We then calculate pairwise cosine similarity between all pathway embeddings to generate a similarity matrix^28,47,49^. Using this matrix, we construct a pathway similarity network, where edges connect pathways with high functional similarity. The community detection algorithms are applied to identify modules within this network: densely connected modules of pathways that represent functional units. These modules, termed functional modules, group together pathways with the same or similar biological functions.

To facilitate the interpretation of these functional modules, namely functional module annotation, MAPA incorporates an LLM-based annotation pipeline (**Methods**). For each functional module, MAPA retrieves related literature from online PubMed linked to its constituent pathways. These resources are provided as input to the LLM to generate a concise functional annotation for the module. The annotation result includes a module title, a functional summary, and, if phenotype information is provided, a context-specific explanation of the module’s relevance to the biological condition. A confidence score is also computed for each annotation to quantify its interpretive reliability.

The final output of MAPA includes both interactive data visualizations and structured functional module reports. Visualization components depict the relationships between molecules, pathways, and functional modules, while module reports summarize all key metadata, including enriched pathways, mapped molecules, annotation details, and module-phenotype associations (**Methods** and **Supplementary Figure 2-3**).

To support broad accessibility, we implemented MAPA in R, the dominant programming language in the bioinformatics community, and provided it in both command-line (R package) and graphical (Shiny App) formats (**Fig. 1d** and **Methods**). Additionally, a Dockerized version enables seamless and easy deployment on local machines, institutional servers, and cloud environments. An online Shiny interface is also available at https://mapashiny.jaspershenlab.com, offering an accessible option for non-programming users. Additionally, MAPA is fully open-source, reinforcing transparency, reproducibility, and extensibility, key priorities in modern omics and bioinformatics research.

### LLM-Based Biological Text Embeddings Accurately Capture Functional Similarity Between Pathways

A central challenge in MAPA is accurately measuring similarity between enriched pathways to identify functional modules. Multiple algorithms exist for measuring pathway similarity^31,32,55^; however, they all suffer from some key limitations (**Supplementary Note**). To overcome these limitations, we developed Biological Text Embedding Similarity (Biotext embedding) (**Methods**). Biotext embedding embeds each pathway into a high-dimensional semantic vector, which can capture complex functional meaning to compare the functional similarity of pathways within and across databases.

To evaluate whether biotext embedding similarity can accurately capture functional similarity both within and across pathway databases, we curated a benchmark pathway dataset, which contains 44 pathways, grouped into 12 biologically meaningful modules based on prior domain knowledge (**Methods**, **Fig. 2a**, and **Supplementary Data 2**). Because conventional semantic similarity approaches like the Wang algorithm are only applicable to GO terms^53^, we first assessed similarity among all 44 entries using only approaches that support cross-database comparisons. We excluded component-based methods with highly redundant correlation profiles (**Supplementary Figure 4**) and retained the Jaccard index and overlap coefficient as representative approaches.

**Figure 2.**
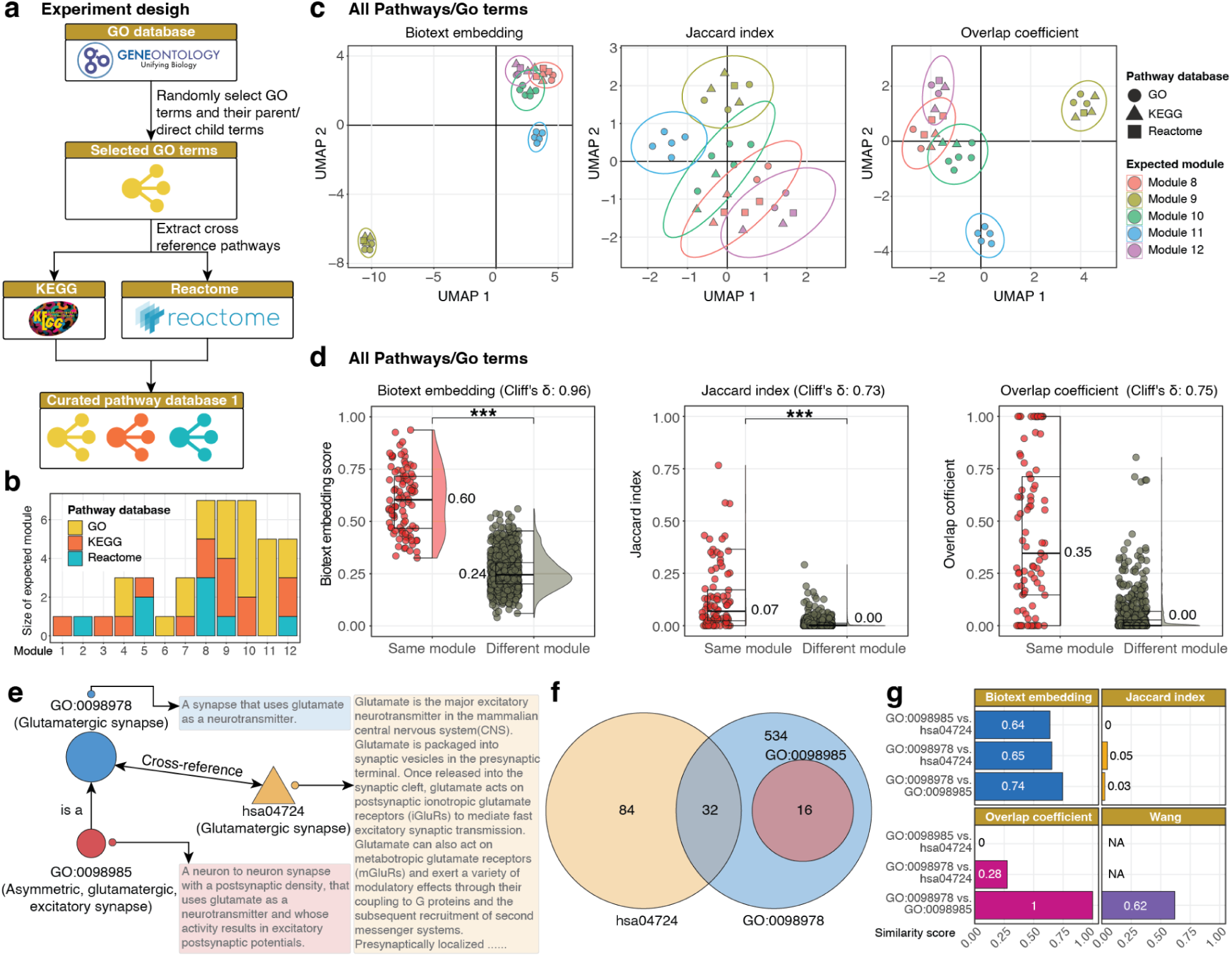
Biotext embedding similarity improves pathway similarity detection within and across databases. (**a**) Overview of the workflow to construct the curated pathway dataset, which includes 44 pathways and GO terms grouped into 12 functional modules. (**b**) Distribution of pathway source across the 12 modules. (**c**) UMAP visualizations of pathway similarity using biotext embedding, Jaccard index, and overlap coefficient. Only modules with ≥ 5 members are shown. (**d**) Comparison of similarity scores between within-module and between-module pathway pairs for all three similarity methods across all databases. (**e**) Case study of one representative module containing three functionally related entries: two GO terms and one KEGG pathway. (**f**) Gene set overlap among the three pathways. (**g**) Pairwise similarity scores among the three pathways using biotext embedding, Jaccard index, overlap coefficient, and Wang (for GO-only pairs). Wilcoxon rank-sum test: *** p-value < 0.001; ** p-value < 0.01; * p-value < 0.05; N.S., not significant.

We computed pairwise similarity matrices using each method and visualized them using UMAP (**Methods** and **Supplementary Data 2**). All three approaches successfully separated five modules (**Fig. 2c**). When comparing similarity scores between within-module and cross-module pathway pairs, the Interquartile Range (IQR) for biotext embedding was 0.47-0.72 for within-module pairs and 0.20-0.30 for cross-module pairs. In contrast, the IQR for the Jaccard index was 0.02-0.17 (within-module) and 0-0.005 (cross-module), while the overlap coefficient yielded 0.16-0.92 (within-module) and 0-0.30 (cross-module) (**Fig. 2d**). Quantifying the discriminative performance, biotext embedding achieved the highest separation between within-module and cross-module similarities (Cliff’s δ = 0.96), outperforming both the overlap coefficient (δ = 0.75) and Jaccard index (δ = 0.73), highlighting its superior capability for distinguishing functionally related pathway pairs. We next evaluated similarity methods specifically for GO terms (**Methods** and **Supplementary Data 2**). Biotext embedding achieved perfect discrimination between within-module and cross-module GO term pairs (δ = 1.0), significantly outperforming traditional methods, including Jaccard index (δ = 0.87) and overlap coefficient (δ = 0.91) (**Supplementary Figure 5**). While Wang semantic similarity matched the biotext embedding in terms of effect size (δ = 1.0), its inability to compute similarities across different GO sub-ontologies resulted in substantial missing data, demonstrating the practical advantage of biotext embedding for cross-ontology functional similarity capture. In summary, these results indicate that biotext embedding more effectively distinguishes biologically coherent functional modules across diverse pathway databases, outperforming conventional similarity metrics.

To further illustrate how different approaches interpret pathway relationships, we examined a representative module containing GO:0098985 (Asymmetric, glutamatergic, excitatory synapse), its parent term GO:0098978 (Glutamatergic synapse), and a KEGG cross-reference, hsa04724 (Glutamatergic synapse) (**Fig. 2e**). Gene content analysis confirmed that GO:0098978 is a subset of GO:0098985, while KEGG hsa04724 shares no overlapping genes with GO:0098985 (**Fig. 2f**). As expected, the Wang method returned a high similarity (0.62) between the GO terms but could not evaluate the KEGG pathway (**Fig. 2g**). In contrast, biotext embedding produced uniformly high similarity scores across all three pathway databases (0.64, 0.65, and 0.74), consistent with their biological connections (**Supplementary Note**).

Together, these findings establish biotext embedding as a robust, generalizable, and database-agnostic similarity metric that captures the intrinsic functional relationships among pathways within and across different databases.

### Functional Module Identification from Pathway Similarity Networks

To cluster biologically related pathways into functional modules, we applied multiple community detection algorithms to the pathway similarity matrix generated using the biotext embedding similarity metric (**Methods**). Functional modules are defined as groups of pathways that share similar or the same biological functions.

We systematically evaluated a range of clustering algorithms commonly employed in biological network analysis (**Methods**). Benchmarking these methods using the curated pathway dataset revealed that graph-based algorithms^56^ slightly outperformed distance-based approaches (**Fig. 3a** and **Methods**). The Binary-cut algorithm^33^ also demonstrated high performance, comparable to graph-based approaches rather than purely distance-based methods. To statistically assess the superiority of graph-based methods, we conducted permutation testing comparing one of the best graph-based methods (Louvain) to the best-performing distance-based method (hierarchical clustering with ward.D2 linkage^57^. The Louvain algorithm significantly outperformed hierarchical clustering (**Fig. 3b**), leading us to adopt Louvain as the default clustering method in MAPA.

**Figure 3.**
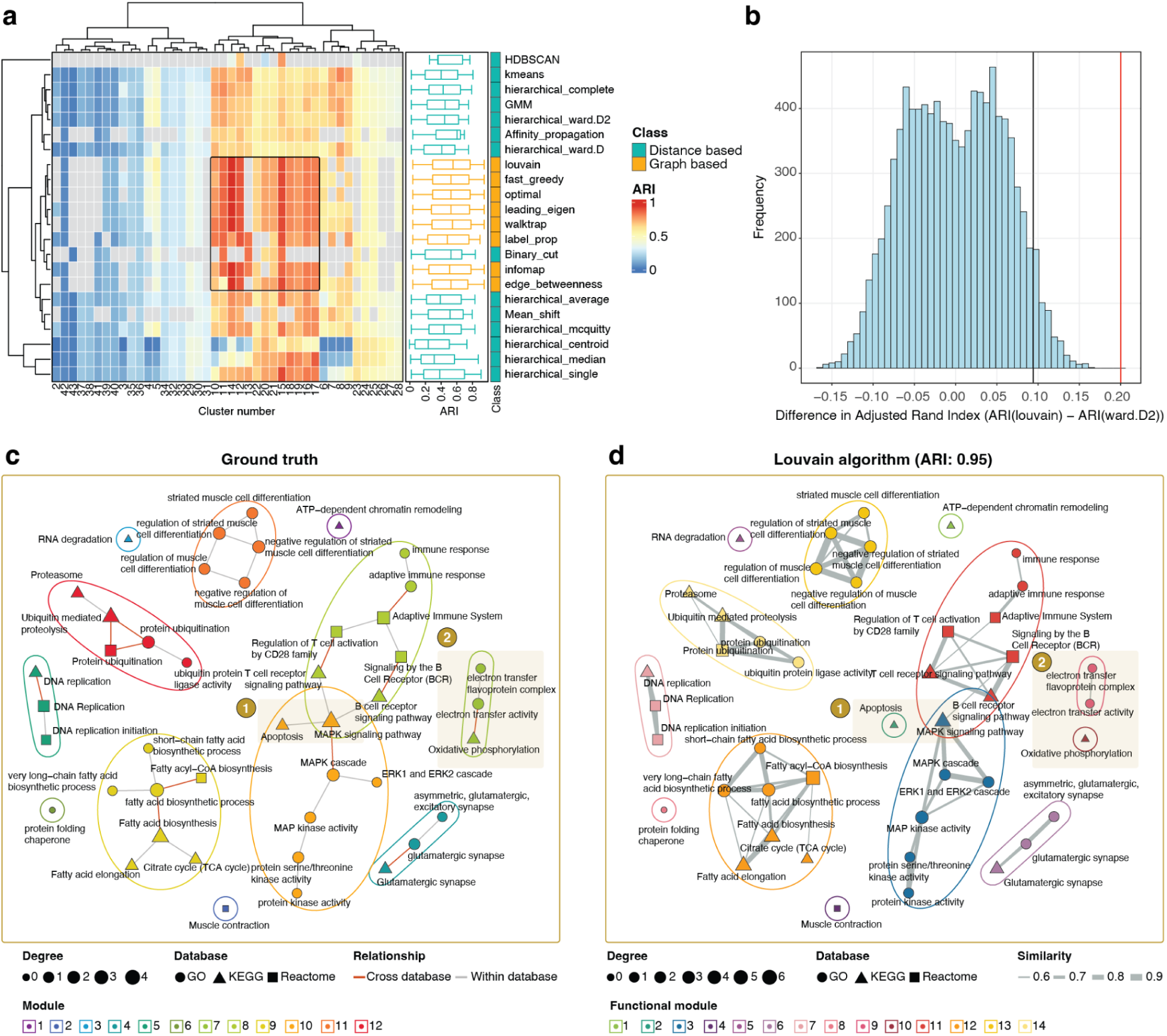
Benchmarking results of functional module identification using multiple clustering approaches. (**a**) Heatmap displaying performance of various clustering algorithms across different parameter settings, measured by ARI relative to ground truth modules. (**b**) Histogram showing permutation test results comparing ARI differences between Louvain and Ward.D2 clustering, confirming Louvain’s superior performance. (**c**) Ground truth network visualization of curated functional modules, with pathways grouped into 12 known biological modules. (**d**) Louvain clustering result showing high concordance with ground truth (ARI = 0.95), with most modules correctly reconstructed.

Further analysis showed that Louvain achieved excellent concordance with the manually curated ground truth, attaining an Adjusted Rand Index (ARI) of 0.95 (**Fig. 3d**). Despite this strong agreement, two slight discrepancies emerged when a similarity cutoff of 0.55 was applied. First, in the ground truth, the Apoptosis pathway (hsa04210) was expected to cluster with pathways involved in kinase signaling, including MAPK signaling, MAPK cascade, ERK1 and ERK2 cascade, MAP kinase activity, protein serine/threonine kinase activity, and protein kinase activity (**Fig. 3c**). However, in the Louvain result, Apoptosis formed a distinct, single-pathway module (**Fig. 3d**). This separation occurred because the biotext embedding generated only moderate similarities (0.33-0.52) between Apoptosis and the other MAPK-related pathways. Since these scores fell below the applied 0.55 similarity cutoff, Apoptosis was isolated as a singleton. The moderate scores are due to the source database descriptions, which emphasize cell survival aspects of MAPK signaling and do not extensively detail its connection to apoptosis^42,58,59^. This highlights the analytical flexibility of the MAPA. A high similarity threshold yields highly specific functional modules, whereas a lower threshold could capture broader themes like overall cell fate control (**Supplementary Note**).

The second difference involved Oxidative phosphorylation (hsa00190), which was grouped in the ground truth with electron transfer activity and electron transfer flavoprotein complex (**Fig. 3c**). In the Louvain clustering, Oxidative phosphorylation was also isolated into its own module (**Fig. 3d**). This was the result of a low similarity score of approximately 0.4 with its expected partners. We hypothesize that this low score is due to the brevity of the pathway’s description (“Oxidative phosphorylation”), which provides insufficient semantic context for the embedding model to form a strong link. This is supported by the observation that other terms with similarly low scores either have very short descriptions or share only superficial words (*e.g.*, “phosphorylation”) with “Oxidative phosphorylation” (**Supplementary Note** and **Supplementary Data 2**).

Collectively, these results demonstrate that Louvain-based community detection accurately reconstructs the major structure of biological functional modules, suggesting that biotext embedding offers finer functional resolution.

### Functional Module Annotation Using LLMs with Retrieval-Augmented Generation

Following the identification of functional modules, a critical next step is to assign biological meaning to each module, a process termed functional annotation. Traditionally, this task has relied on heuristic or manual methods^26–28,33,36,60,61^. Some methods annotate functions of modules by word frequencies of the pathway names^36,61^, while others use the pathway with the smallest enrichment *p-value* or the one with the highest average similarity to other members of the module^26–28^. While convenient, these strategies are inherently biased and often fail to reflect the collective biological significance of the entire module.

To this end, we developed a robust pipeline for LLM-based annotation of functional modules (**Fig. 4a** and **Methods**). For each functional module, we first extract comprehensive information on the constituent pathways from standard databases. Next, to contextualize the module, we retrieve related literature from PubMed and database-linked references. Additionally, users can upload relevant literature manually, ensuring coverage of unpublished or non-indexed research. Then, similarities are calculated between the modules and each literature to identify the top 20 most relevant literatures. These are further re-ranked by the LLM, which selects the top 5 publications that best match the functional profile of the module (**Methods**).

**Figure 4.**
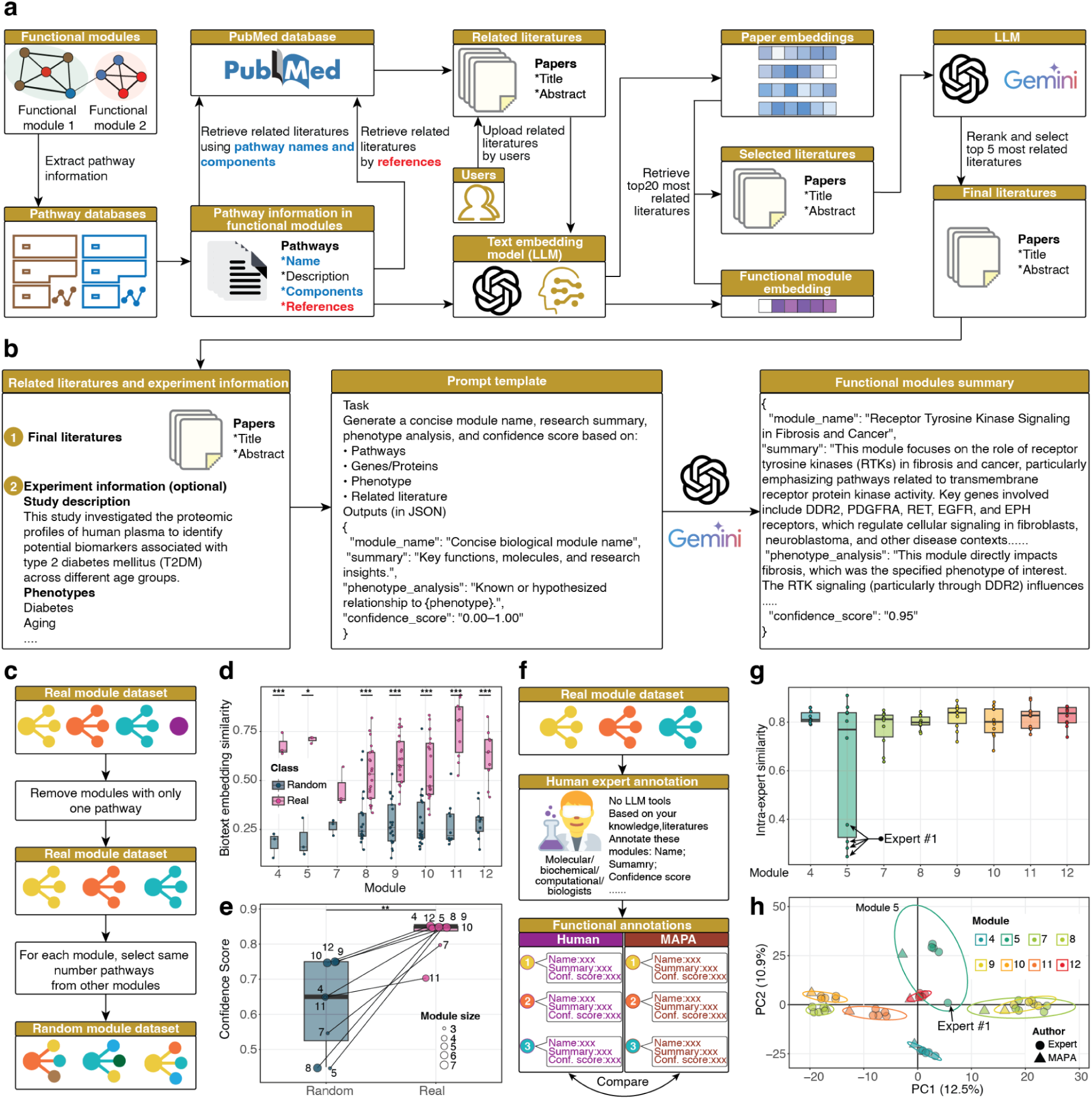
Functional module annotation using LLMs with RAG and evaluation of its performance. (**a**) Overview of the functional annotation pipeline in MAPA. (**b**) Final functional annotation step. The selected literature, along with optional user-provided experimental or phenotype context, is fed into a structured prompt template guiding the LLM to generate a comprehensive annotation for functional modules. (**c**) Schematic workflow for generating a negative control (random module) dataset. (**d**) Boxplots comparing intra-module biotext embedding similarity scores between real and random module datasets. (**e**) Comparison of confidence scores between real and random functional modules. (**f**) Workflow for human expert annotation. Experts reviewed real functional modules and generated annotations, which were then compared with MAPA-derived annotations to assess consistency and quality. (**g**) Intra-expert similarity of functional annotations for each module. For most modules, human experts produced highly consistent annotations, except for module 5, where expert #1 provided a markedly divergent interpretation. (**h**) PCA of functional annotations from human experts and MAPA. Annotations from MAPA (triangles) cluster closely with those from human experts (circles) across all modules, except module 5, where expert #1’s annotation deviates substantially from both MAPA and other experts. Wilcoxon rank-sum test: *** p-value < 0.001; ** p-value < 0.01; * p-value < 0.05; N.S., not significant.

In the final annotation step, we combine the curated module content, literature set, and optionally, user-provided phenotype descriptions from the original experiment (**Fig. 4b**). This information is fed into a carefully designed prompt instructing the LLM to generate a comprehensive annotation (**Supplementary Note**). The prompt asks the model to produce: (1) a concise title for the functional module, (2) a biological summary integrating the module’s purpose and mechanisms, (3) an analysis of its potential relationship to the phenotype under study, and (4) a numerical confidence score reflecting how strongly the molecular and pathway content converges on a coherent biological process (**Fig. 4b**).

### Performance Evaluation of MAPA’s Functional Annotation

To rigorously evaluate the performance of functional annotation by MAPA, we conducted a negative control experiment using a randomly constructed pathway dataset (**Fig. 4c** and **Methods**). Starting from the curated pathway dataset, we retained only modules containing at least two pathways. For each real module, we then generated a random module by sampling the same number of pathways from the pool of all other pathways outside the original module (**Supplementary Note**). This process ensured that random modules matched the real modules in size but lacked any underlying biological coherence.

We first compared the intra-module biotext embedding similarity scores between real and random modules. As expected, real modules exhibited significantly higher intra-module similarity than random modules (*p-values* = 0.0006, 0.0103, 0.0637, 1.3 × 10^−6^, 1.3 × 10^−10^, 1.4 × 10^−6^, 2.0 × 10^−7^ and 5.0 × 10^−6^; Wilcoxon rank-sum test), indicating that pathways within true functional modules are more functionally related than those grouped at random (**Fig. 4d**). This also confirmed that random modules lack meaningful biological structure. Next, we applied our LLM-based annotation pipeline to both real and random modules (**Methods**). The confidence scores for random modules were significantly lower than those for real modules (*p-value* < 0.031; Wilcoxon rank-sum test; **Fig. 4e**), demonstrating that the scoring system effectively discriminates biologically meaningful modules from arbitrary groupings.

To further validate the biological relevance and reliability of MAPA’s annotations, we conducted a human expert comparison experiment involving five specialists in molecular biology, biochemistry, and computational biology (**Fig. 4f** and **Methods**). Each expert was provided with the names and descriptions of pathways within real functional modules and instructed to generate annotations equivalent in scope and detail to those produced by the MAPA pipeline (**Supplementary Table 2**). We first assessed the consistency of annotations across experts. All textual summaries were converted into text embeddings, and intra-expert similarity for each module was calculated using cosine similarity (**Methods**). As anticipated, most modules demonstrated strong agreement among experts, with mean similarity values exceeding 0.8 (**Fig. 4g**). The only exception was module 5, where Expert #1 provided annotations that differed substantially from the others (**Fig. 4g**). Upon review, Expert #1 had entered “Not Available” for both the module name and summary (**Supplementary Table 2**). This discrepancy explains the observed deviation for module 5 and highlights the variability introduced by human judgment in annotation tasks.

Next, we evaluated the concordance between MAPA’s annotations and those generated by human experts. All annotations, both human expert- and MAPA-generated, were converted into embeddings and visualized using PCA (**Methods**). For nearly all modules, MAPA’s annotations clustered tightly with those of the experts, demonstrating strong alignment between the MAPA and expert-driven interpretations (**Fig. 4h**). The only outlier was module 5, which, as noted, reflects the incomplete annotation provided by Expert #1 rather than a limitation of MAPA (**Supplementary Table 2**). These results confirm that MAPA delivers annotation quality comparable to experienced domain experts while maintaining consistency and avoiding errors stemming from subjective interpretation.

Finally, we tested MAPA’s reproducibility of annotation across 15 independent runs on the curated pathway dataset (**Methods**). The clustering step was perfectly stable, yielding 14 identical modules each time (pairwise ARI = 1). For the generated modules with more than one pathway, LLM-generated annotations showed high consistency for the same module across runs (cosine similarity > 0.90), contrasting sharply with the low similarity between different modules. This confirms that MAPA reliably produces stable and specific functional interpretations (**Supplementary Figure 7**).

Collectively, these results demonstrate that MAPA’s functional annotation pipeline accurately captures biologically meaningful insights and produces expert-level interpretability at scale. Unlike human manual annotation, which is time-consuming, subjective, and prone to occasional inconsistencies, MAPA provides an automated and reproducible solution for generating reliable module-level interpretations.

### Benchmarking MAPA Against Existing Pathway Redundancy Reduction Approaches

We next sought to benchmark MAPA’s performance against several widely used and representative tools. Specifically, we compared MAPA to PAVER^28^, enrichplot^36^, and aPEAR^26^, all of which are commonly used and typical tools designed to address pathway redundancy in pathway enrichment analyses.

We first evaluated whether these tools could accurately identify true functional modules using the curated pathway dataset (**Fig. 5a** and **Methods**). As expected, MAPA achieved the highest performance with an ARI of 0.95, substantially exceeding the ARIs of aPEAR (0.33), enrichplot (0.33), and PAVER (0.23), indicating superior accuracy in reconstructing biologically coherent modules (**Fig. 5b, Supplementary Data 2,** and **Supplementary Figure 6**). The functional modules identified by MAPA showed a clear, near one-to-one correspondence with the ground truth clusters, whereas all other approaches exhibited substantial discrepancies, frequently failing to group related pathways into the same module (**Fig. 5b**).

**Figure 5.**
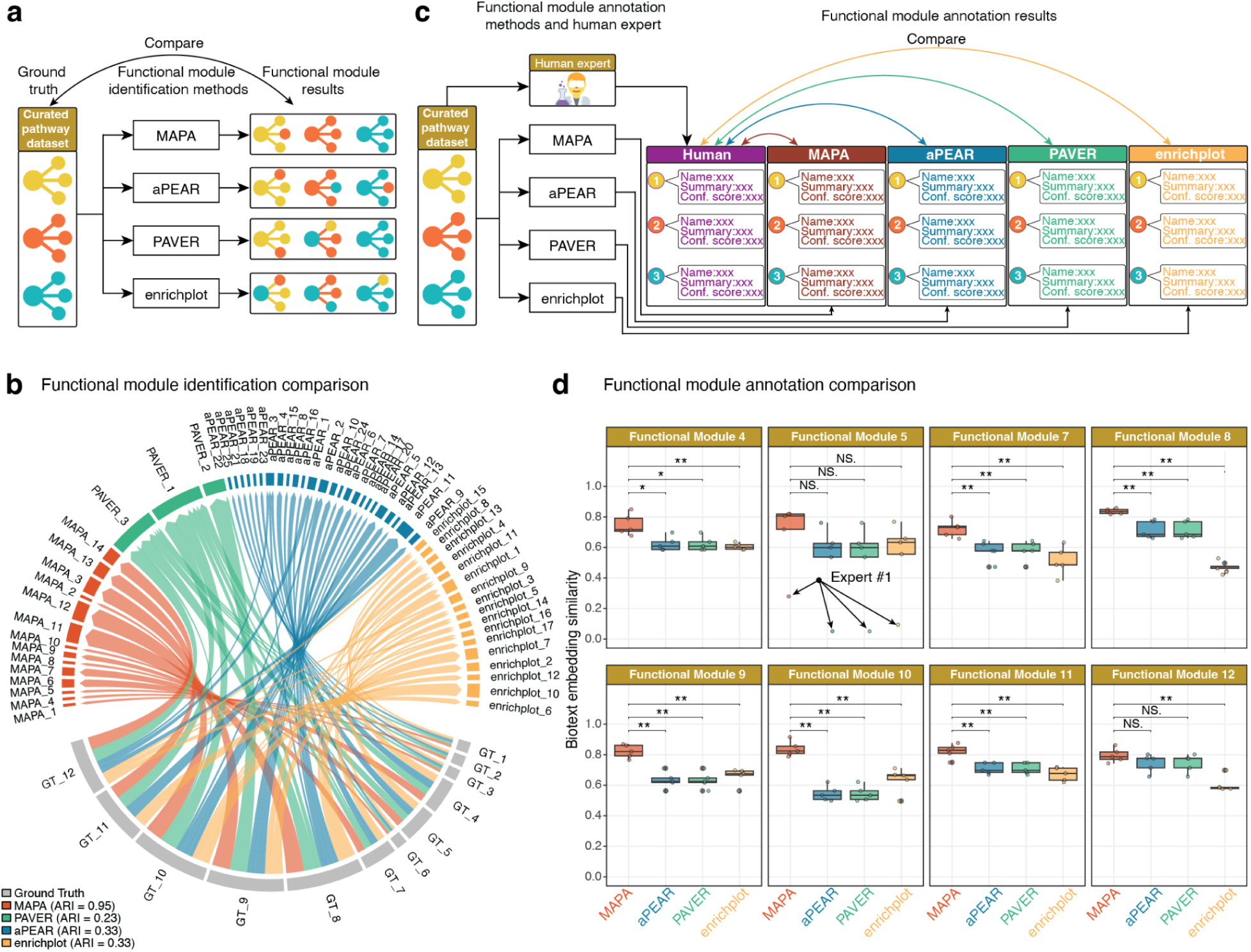
Benchmarking MAPA against existing pathway redundancy reduction approaches. **(a)** Schematic workflow for benchmarking functional module identification between MAPA and other approaches. **(b)** Benchmarking module identification against ground truth. The circos plot illustrates the mapping of pathways from modules identified by different methods to the ground truth modules. The outer segments represent the modules, and connecting ribbons show the cluster assignments. The near one-to-one mapping between MAPA (red) and the ground truth (grey) demonstrates high concordance, in contrast to the over-clustering by PAVER (green) and fragmentation by aPEAR (blue) and enrichplot (orange). **(c)** Schematic workflow for benchmarking functional module annotation between MAPA and other approaches. **(d)** Box plots comparing the cosine similarity between expert-curated annotations and those generated by MAPA, aPEAR, PAVER, and enrichplot. Across most functional modules, MAPA’s annotations demonstrate a significantly higher similarity to the expert standard. Wilcoxon rank-sum test: *** p-value < 0.001; ** p-value < 0.01; * p-value < 0.05; N.S., not significant.

Next, we further assessed the ability of each tool to generate accurate functional annotations (**Fig. 5c** and **Methods**). Eight real modules with expert-curated annotations (**Fig. 4f**) were used as a reference standard. Each method was applied to annotate these modules, and the semantic similarity between the tool-generated annotations (**Supplementary Data 2**) and those from five human experts was calculated. MAPA consistently outperformed all other methods, producing annotations that were significantly closer to expert interpretations (Wilcoxon rank-sum test) (**Fig. 5d**). The only exception occurred in Module 5, where one expert provided an invalid annotation (“Not available”), explaining the observed variance (**Supplementary Table 2**).

Collectively, these results demonstrate that MAPA not only delivers superior accuracy in functional module identification but also provides more informative and biologically meaningful annotations compared to existing approaches. Detailed benchmarking metrics and comparisons are summarized in **Supplementary Table 1.**

### MAPA Reveals Comprehensive Multi-Organ Functional Dysregulation in Aging

To demonstrate MAPA’s utility in a real-world biological context, we applied it to a comprehensive monkey multi-organ aging dataset across two age groups (3-5 and 16-19 years) representing young and old populations^3^. This dataset encompasses multi-omics profiles, which provide a rich multi-omics view of aging across the organism (**Methods**). The age-dependent molecules were identified by fuzzy clustering. For this analysis, we focused on the clusters that were continuously upregulated or downregulated during aging, and applied MAPA to each omics layer (**Supplementary** Figures 8-13).

Specifically, for transcriptomics data, MAPA identified 162 distinct functional modules from 463 unique enriched pathways across 56 tissues (**Supplementary Data 3** and **Supplementary** Figures 12-13). For example, consistent with the pathway enrichment results, we also found that the inflammatory response relevant functional module “Chemokine signaling in inflammation” increased in lung (LILU) tissue, and lipid metabolism relevant functional module “AMPK-mTOR pathway in metabolic control” is decreased in skin (neck) tissue, which are consistent with the original paper^3^.

To enhance interpretability and focus on robust patterns, we retained only the dysregulated functional modules that were observed in more than 3 tissues, and further limited our interpretation to tissues that exhibited at least 2 functional modules. For each functional module in each tissue, we also calculated the dysregulation index to reflect their changes in aging (**Methods**) (**Fig. 6a**). Some interesting results were found. For example, distinct brain regions exhibited coordinated age-associated changes. Specifically, the frontal lobe (FL) and parietal lobe (PL) clustered together (**Fig. 6a**), characterized by an upregulation of modules related to neurodegenerative pathways across tissues and multi-tissue autophagy regulation, reflecting common neurodegenerative and proteostatic responses during aging (**Fig. 6b**). Moreover, we observed cross-organ clustering along the brain-gut axis: the duodenum and brain (FL) jointly showed upregulation of iron metabolism and cell death pathways, suggesting shared vulnerability to ferroptosis-related processes (**Supplementary Figure 8a**). Similarly, the duodenum and brain (PL) clustered together (**Fig. 6a**), marked by upregulation of proteasomal regulation in multi-tissue biology and RNA processing and mitochondrial function in disease (**Fig. 6c**), pointing to coordinated dysregulation of protein homeostasis and mitochondrial gene expression across neural and intestinal tissues.

**Figure 6.**
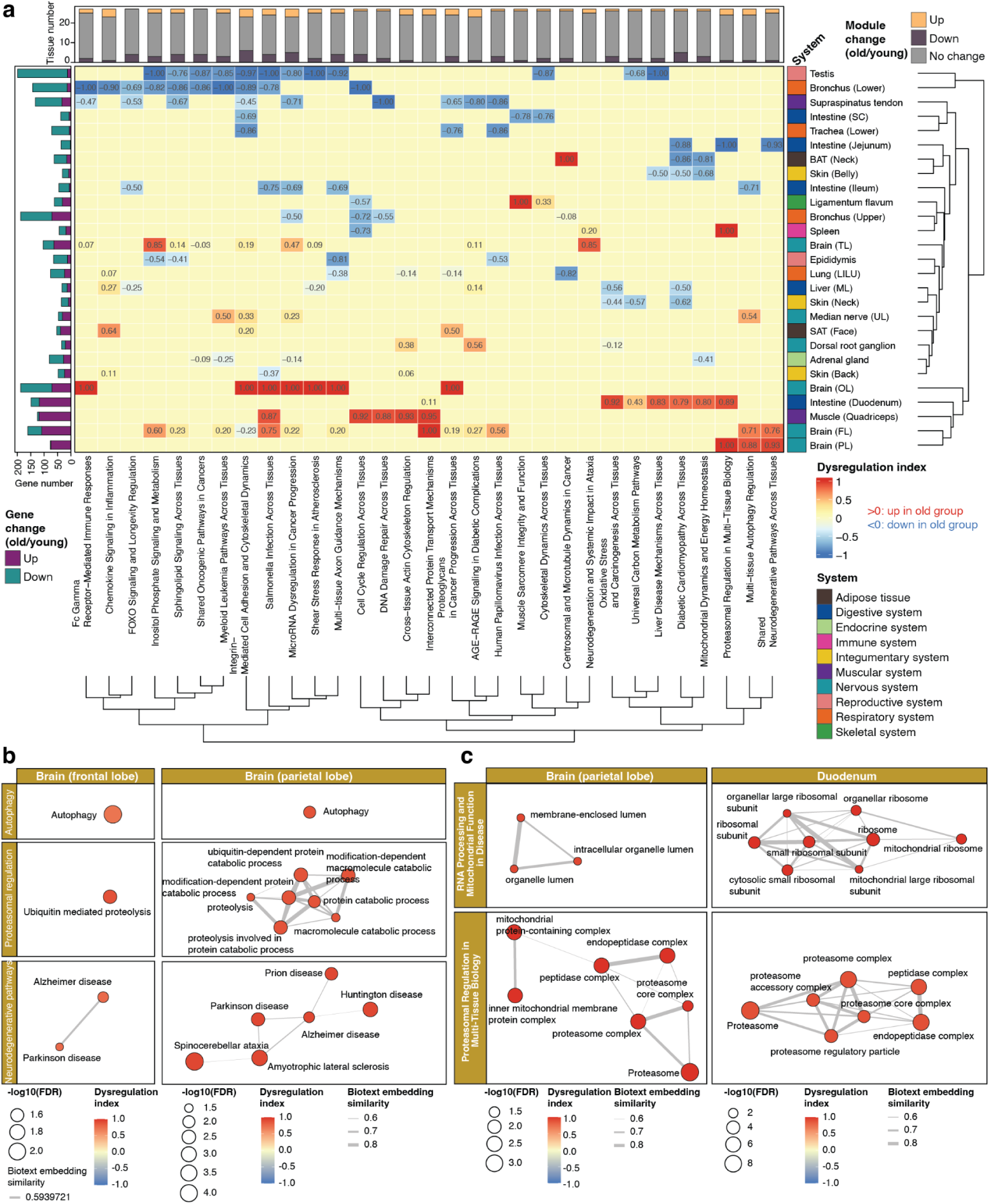
MAPA reveals functional dysregulation across multiple organs during aging. **(a)** Heatmap of dysregulation index for selected functional modules (rows) across aging tissues (columns), calculated from transcriptomic data using MAPA. Red indicates upregulation in the old group; blue indicates downregulation. **(b)** MAPA uncovered coordinated activation of neurodegeneration and proteostasis-related modules, such as autophagy, ubiquitin-mediated proteolysis, and Alzheimer’s disease, in both frontal and parietal lobes. **(c)** Shared upregulated functional modules between brain and gut tissues highlight cross-organ aging programs. The parietal lobe and duodenum both show coordinated upregulation of proteasome-related and mitochondrial function modules.

In contrast, we observed consistent downregulation of FOXO signaling and longevity regulation across diverse tissues, including the bronchus (lower), supraspinatus tendon, ileum, and liver (median lobe), highlighting a conserved decline of longevity-associated transcriptional programs during aging (**Supplementary Figure 8b**). Additionally, microRNA dysregulation in cancer progression emerged as a cross-tissue signature with both up- and down-regulation across nine tissues (**Fig. 6a**). Notably, this module was downregulated in the testis, bronchus (upper), ileum, supraspinatus tendon, and adrenal gland, whereas it was upregulated in several neural tissues, including the frontal and occipital lobes, temporal lobe, and median nerve (ulnar) (**Supplementary Data 3**). This bidirectional regulation pattern suggests tissue-specific trajectories of microRNA-mediated control during aging, with suppression in peripheral and barrier tissues but activation in neural compartments.

For the plasma proteomics data, MAPA provided deeper functional insights than the original study. For example, from the list of downregulated proteins (cluster D), MAPA identified a functional module named “Autophagic and Endosomal Pathways in Cellular Homeostasis.” (**Supplementary Data 3** and **Supplementary Figure 11a**) This result refines the output from the original paper, which only identified “Autophagy” as a significant pathway. By clustering related pathways, MAPA captured the critical interplay between autophagy and endosomal transport. The LLM-generated summary aligns with the broader understanding that impaired autophagy is a key contributor to aging and age-related diseases^62^. Notably, the annotation highlighted key players, including Vacuolar Protein Sorting (VPS) proteins whose dysfunction in trafficking autophagy proteins is directly implicated in neurodegenerative conditions like Parkinson’s disease^63^.

Together, these results illustrate MAPA’s power to reveal biologically meaningful, multi-organ aging signatures by resolving redundancy and fragmentation in conventional pathway analyses. MAPA offers a robust framework for dissecting complex aging processes at the systems level.

## Discussion

In this study, we developed MAPA, an innovative and comprehensive computational framework designed to systematically resolve redundancy in pathway enrichment analyses and enhance the biological interpretability of multi-omics data. MAPA introduces several significant advances over existing tools for pathway analysis and redundancy resolution. First, MAPA leverages LLM-based biotext embeddings to quantify functional similarity between pathways across different and latest databases. By embedding pathway names and functional descriptive texts into semantic vectors, MAPA captures nuanced functional relationships that extend far beyond simple molecule overlap or rigid ontology hierarchies. This allows it to identify functional relationships between pathways even when they share few or no molecular components, bridging gaps that conventional methods cannot address. Second, MAPA is capable of analyzing pathway enrichment results from various pathway databases. Unlike many redundancy-reduction methods that are limited to the GO or specific resources, MAPA’s biotext embedding-based similarity framework is database-agnostic. This flexibility facilitates the integration and comparison of pathways from diverse knowledge bases, a critical capability as biological databases become increasingly interconnected and heterogeneous. Third, MAPA advances automated, literature-informed functional module annotation based on LLMs. Existing tools typically select a single representative pathway to label each module, potentially losing the broader biological context, which can result in biased interpretations. In contrast, MAPA synthesizes information from all pathways within a module, supplements it with relevant scientific literature retrieved through automated searches, and generates comprehensive, human-readable annotations. This method not only improves the interpretability of results but also provides users with richer biological narratives that reflect the collective function of the module rather than isolated pathways. Finally, MAPA supports broad multi-omics applications, encompassing transcriptomics, proteomics, and metabolomics data. This versatility allows researchers to apply a consistent, redundancy-aware analytical framework across multiple data modalities, facilitating integrated insights into complex biological processes. Together, these advances position MAPA as a transformative tool for multi-omics data analysis, enabling researchers to move beyond lists of enriched pathways toward deeper, redundancy-resolved biological interpretation.

Benchmarking analyses demonstrate that MAPA outperforms established redundancy-reduction tools^26,28,36^. MAPA achieves higher accuracy in reconstructing known functional modules (module identification), as evidenced by its higher ARI scores in curated benchmark datasets. Moreover, MAPA’s capability for high-quality, literature-backed annotations distinctly surpasses tools focusing solely on clustering or visualization (module annotation).

The application of MAPA to the multi-organ monkey aging dataset provided new biological insights into the functional dynamics of aging across diverse tissues^3^. Unlike conventional pathway analysis methods that often yield lengthy and redundant lists of pathways, MAPA distilled these results into coherent functional modules, allowing for more interpretable and biologically meaningful patterns. This approach enabled the identification of novel cross-tissue regulatory programs, such as multi-tissue autophagy regulation, AMPK-mTOR signaling in metabolic control, and RNA processing and mitochondrial function in disease, which are hidden by conventional pathway analysis methods. MAPA’s tissue clustering based on functional module dysregulation also uncovered biologically coherent groupings, such as coordinated aging responses across brain regions, and gut-brain associations that invite further mechanistic exploration. These findings suggest that aging induces not only organ-specific but also system-wide shifts that may reflect shared metabolic, proteostatic, or inflammatory burdens. Altogether, this analysis demonstrates MAPA’s ability to uncover biologically plausible and hypothesis-generating insights from complex multi-omics datasets. By resolving pathway redundancy and contextualizing functional changes at the tissue level, MAPA enables a systems-level view of aging and offers a valuable framework for future mechanistic investigations and biomarker discovery.

While MAPA offers substantial advances in redundancy-aware pathway analysis and functional module interpretation, it also presents opportunities for future development. Currently, MAPA analyzes individual omics datasets separately and does not yet integrate multiple omics layers into a unified analysis pipeline, potentially overlooking coordinated molecular interactions and cross-regulatory mechanisms spanning different biological levels. Future extensions aim to integrate multiple omic layers to identify cross-omics functional modules that reflect the coordinated regulation of complex biological processes. Additionally, the computational demands of LLM use present a challenge. Generating high-dimensional text embeddings and performing LLM-based annotation, particularly at scale, requires access to commercial LLM APIs and may involve significant costs or require specialized hardware or cloud computing resources. Future development plans include exploring open-source LLM alternatives such as Llama^64^, Qwen^65^, and DeepSeek^66^, and developing strategies for local deployment of these models. In addition, optimization techniques like embedding caching, quantization, and model distillation could substantially reduce computational requirements without compromising performance.

MAPA exemplifies the transformative potential of LLMs and advanced machine learning in the evolving landscape of multi-omics and computational biology. As omics technologies continue to generate ever-larger and more complex datasets, there is an increasing demand for tools that can not only handle high-dimensional data but also provide deep biological interpretation that transcends simple statistical associations. By leveraging LLM-based embeddings and functional annotation, MAPA offers a blueprint for how AI can be integrated into bioinformatics workflows to deliver insights that would previously have required extensive manual curation or domain expertise. Looking forward, MAPA exemplifies the broader vision of integrating LLM-based tools into routine bioinformatics pipelines. As LLM technologies continue to advance, we anticipate that tools like MAPA will become standard components of omics analysis, enabling researchers to explore complex biological questions with unprecedented speed, depth, and interpretive clarity. Ultimately, such innovations promise to accelerate our understanding of health and disease and to facilitate the development of more precise, personalized therapeutic strategies.

## Methods

### Construction of curated pathway datasets

Two curated pathway datasets were constructed to demonstrate the application of MAPA and benchmarking.

#### Curated pathway dataset 1

To construct the curated pathway dataset 1, we aimed to create a robust ground truth for evaluating our method’s ability to cluster functionally related pathways and distinguish unrelated ones. We began by identifying functionally unrelated GO terms from the AmiGO2 database (https://amigo.geneontology.org/amigo/dd_browse), which served as seeds for our expected functional modules. For each selected GO term, we consulted QuickGO (https://www.ebi.ac.uk/QuickGO/) to gain a comprehensive view. We then expanded these initial seeds into expected functional modules by retrieving their immediate “is a” or “part of” parent and child terms, and any cross-reference (xref) pathways recorded in KEGG or Reactome. When a seed lacked an xref entry, we performed a manual name-based search in KEGG and Reactome to capture pathways bearing an identical or synonymous name. Additionally, to ensure a clear separation of unrelated functions, we specifically selected four pathways from GO, KEGG, and Reactome that appeared functionally distinct from any of our established modules, designating them as singletons. This rigorous curation ensured that our dataset comprised both tightly coherent functional groups and clearly isolated unrelated pathways, providing a reliable benchmark for our clustering methodology. The final curated pathway dataset 1 comprised 12 expected functional modules: 8 multi-term modules, each containing 3 to 7 pathways, and 4 singletons. In total, this dataset encompassed 44 distinct pathway records, distributed across 22 GO terms, 14 KEGG pathways, and 8 Reactome pathways. This matrix constitutes the ground truth against which MAPA’s clustering performance is assessed.

#### Curated pathway dataset 2

Curated pathway dataset 2 centers on metabolic functions. Our objective was to demonstrate the substantial redundancy of metabolic pathways across major public pathway databases. To this end, we applied the Jaro-Winkler similarity metric to perform fuzzy matching of pathway names across SMPDB, KEGG, and Reactome, setting a similarity cutoff of 0.85 to balance tolerance to minor lexical variation against the risk of false positives. Unmatched pathways were subsequently refined through manual curation to enhance the reliability of the matched set. The above procedures resulted in 44 matched pathway pairs between Reactome and KEGG, 43 between SMPDB and Reactome, and 44 between SMPDB and KEGG. These matched pathways were compiled into the curated pathway dataset 2 (**Supplementary Data 2**).

### Sample preparation and analytical conditions for the case study

This dataset is from a previously published study^3^ investigating aging and metformin treatment effects in male cynomolgus monkeys across different age groups (3-5, 10-12, and 16-19 years). The detailed sample preparation and analysis conditions could be found in the previous study^3^. Briefly, the multi-omics data includes bulk RNA sequencing data from 79 tissues using the MGI DNBSEQ-T7 platform, plasma proteomics data using Data Independent Acquisition (DIA) on Thermo Scientific Q Exactive HF mass spectrometer, and plasma metabolomics data using UPLC-MS/MS on ACQUITY 2D UPLC system paired with Q Exactive mass spectrometer. Age-dependent molecular changes were identified using fuzzy c-means clustering (*Mfuzz* R package), revealing four distinct temporal patterns, including continuous upregulation (Cluster U) and continuous downregulation (Cluster D) with aging. For our analysis, we utilized the age-dependent genes, proteins, and metabolites from Cluster U and Cluster D as identified in the original publication^3^.

### Design of the MAPA

We developed MAPA using the R programming language, which is extensively used in bioinformatics and multi-omics research due to its rich ecosystem of statistical and data analysis packages. The core design philosophy of MAPA is to provide a flexible, interpretable, and modular framework for resolving functional redundancy in pathway enrichment results.

MAPA is structured to guide users through a comprehensive workflow, from pathway enrichment and semantic similarity computation to the identification and annotation of functional modules using LLMs. Each component of MAPA is designed to be both standalone and interoperable, allowing seamless integration into custom pipelines or use as a complete end-to-end solution.

### LLMs integration and prompt-based annotation in MAPA

MAPA integrates LLMs at two critical stages: (1) biotext embedding and (2) functional module annotation. For biotext embedding, MAPA supports OpenAI’s embedding models (default: text-embedding-3-small), Google Gemini models (default: models/gemini-embedding-exp-03-07), and SiliconFlow models (default: Qwen/Qwen3-Embedding-8B). These models transform pathway names and descriptions into high-dimensional vector representations that capture the semantic meaning of each pathway.

For functional module annotation, MAPA supports OpenAI’s GPT models (default: gpt-4o-mini-2024-07-18), Google Gemini models (default: gemini-2.5-pro), and SiliconFlow models (default: Qwen3/Qwen3-32B). Users can select any of these platforms for annotation based on their access and preferences. Authentication and setup details can be found in the official documentation. OpenAI API: https://platform.openai.com/api-keys, and Google Gemini API: https://ai.google.dev/gemini-api/docs. MAPA includes built-in functions to streamline API usage and manage requests, enabling seamless and reproducible integration of LLMs into pathway analysis workflows.

The application of LLMs to functional module annotation in MAPA is supported by numerous studies showing that general-purpose language models can solve complex domain-specific tasks through prompt learning alone^47,49^. Prior studies have demonstrated that LLMs, without requiring supervised training in the target task, can accurately interpret biomedical contexts and even perform quantitative predictions, purely from prompt tuning^48^.

This capability arises from the large-scale pretraining process, during which LLMs are exposed to extensive biomedical literature (*e.g.*, PubMed abstracts, pathway databases, and clinical documents). By predicting next tokens in this diverse corpus, the model learns latent representations that encode statistical relationships between biological entities, co-occurrence patterns of molecular functions, and associations among disease terms, pathways, and molecular processes. When prompted with structured inputs describing metabolite sets or pathway modules, these pretrained representations are activated, enabling the model to infer relevant biological functions based on proximity to known semantic clusters (*e.g.*, oxidative stress, immune activation, and lipid metabolism).

In MAPA, we design targeted prompts to elicit such reasoning behavior from the model. The resulting annotations and scores reflect probabilistic mappings within the model’s latent space, rather than keyword matching or rule-based logic. This approach ensures both interpretability and adaptability across pathway types and biological contexts.

### Prompt engineering in MAPA

Two steps in MAPA leverage prompts to instruct the LLM to complete the tasks.

#### Related literature re-ranking

To further refine literature selection beyond embedding-based similarity, MAPA employs a two-stage reranking strategy incorporating LLMs with structured prompts tailored for biomedical relevance assessment (**Fig. 4b** and **Supplementary Note**). The prompt is designed with five key components. (1) Task description: The LLM is instructed to act as a biomedical literature analyst tasked with evaluating whether a given abstract or text chunk is relevant to a functional module defined by specific pathways and molecules (genes, proteins, or metabolites). (2) Module context: For each module, MAPA provides a concise textual summary containing enriched pathway names and the associated molecular entities. This gives the LLM a biological grounding to assess alignment between the module and candidate literature. (3) Evaluation criteria: The prompt guides the model to consider biological relevance (*e.g.*, mechanistic alignment with the module), to disregard superficial mentions or irrelevant metadata (*e.g.*, author names or affiliations), and to provide both a cleaned text and a relevance score ranging from 0.00 to 1.00. (4) Example response format: A structured output format is included in the prompt, instructing the model to return a JSON-style response with two fields: relevance_score and cleaned_text. This ensures consistency and facilitates downstream filtering. This design allows MAPA to retrieve the literature most relevant to the functional module, thereby ensuring the accuracy and reliability of the functional annotation.

#### Functional module annotation

To enable high-quality, automated interpretation of functional modules, MAPA uses LLMs with a structured prompt designed for biological summarization and annotation (**Fig. 4b** and **Supplementary Note**). This prompt consists of six key components. (1) Task description: The prompt opens by assigning the LLM the role of a “molecular and biochemical biologist” and clearly defining the task, to generate a concise biological module name, a comprehensive research summary, a phenotype-specific analysis, and a numerical confidence score. (2) Detailed task instructions: These guide the LLM to identify the core biological process underlying the module, synthesize relationships among pathways and molecules, analyze relevance to a provided phenotype (if applicable), and calculate a confidence score (ranging from 0.00 to 1.00) that reflects the functional coherence of the module. (3) Example input: A representative input is included, featuring example pathways, related genes/proteins, phenotype of interest, and literature abstracts. This helps the LLM learn the structure and expectations of the task. (4) Example output: A formatted JSON response is provided as a template, showing the expected style and depth of output across the four requested fields: module_name, summary, phenotype_analysis, and confidence_score. (5) Actual input: For real tasks, MAPA compiles enriched pathways and associated molecules within each module, retrieves supporting literature through PubMed search or pathway database references, and assembles these into the prompt. If available, the user-provided phenotype description is also included. (6) Final output: The LLM produces a structured JSON response containing a meaningful module title, an integrated biological summary, a phenotype relevance interpretation, and a confidence score. This design enables MAPA to perform rich, literature-informed annotation of functional modules with a consistent format, offering greater interpretability than conventional single-pathway-based summaries.

### Input files for MAPA

Users are required to provide a list of molecular markers as input for MAPA. These markers may be derived from transcriptomics, proteomics, or metabolomics studies, and should consist of gene, protein, or metabolite identifiers, respectively. The format of the input depends on the type of pathway enrichment analysis being performed. Specifically, MAPA accepts either: (1) A list of differentially expressed molecules for Overrepresentation Analysis (ORA), or (2) a ranked list of molecules for Gene Set Enrichment Analysis (GSEA). Currently, GSEA is supported only for transcriptomics and proteomics data. For metabolomics datasets, MAPA supports ORA-based enrichment only, due to limitations in rank-based analysis for small and sparse metabolite lists. For the gene or protein, the user needs to provide at least Ensembl gene IDs (*e.g.*, “ENSG00000141510”), NCBI Entrez gene IDs (*e.g.*, “7157”), UniProtKB accession numbers (*e.g.*, “P04637”), or Gene symbols (*e.g.*, “TP53”). For the metabolite, the user needs to provide either the HMDB ID or the KEGG ID.

### Pathway enrichment analysis

MAPA supports two widely used methods for pathway analysis: ORA^54^ and GSEA^8^. ORA is available for transcriptomics, proteomics, and metabolomics data, while GSEA is currently supported only for transcriptomics and proteomics data due to methodological limitations in rank-based analysis for metabolite datasets.

#### ORA

For transcriptomics and proteomics datasets, MAPA performs ORA using the clusterProfiler R package^67^. By default, three major pathway databases are supported: (1) Gene Ontology (GO) (Biological Process, Molecular Function, and Cellular Component), (2) KEGG, and (3) Reactome. Users can choose one or more of these databases. The default significance threshold for adjusted *p-values* is 0.05, but this can be modified by the user.

For metabolomics data, MAPA leverages the metpath package from the TidyMass project to conduct enrichment analysis^68–70^. The default pathway databases include: (1) KEGG metabolic pathways^42^, and (2) SMPDB (Small Molecule Pathway Database)^45^. As with transcriptomics and proteomics, users can customize the choice of databases. ORA results are filtered using a default adjusted *p-value* cutoff of 0.05. The molecular identifiers used for metabolomics (*e.g.*, KEGG Compound IDs, HMDB IDs) are automatically matched to the selected database using built-in mapping utilities.

#### GSEA

For transcriptomics and proteomics, GSEA is implemented using the clusterProfiler package^67^ as well. This method requires a ranked list of genes or proteins, typically ordered by a metric such as log fold change or a statistical score (*e.g.*, t-statistic, signal-to-noise ratio, or correlation). The same three pathway databases, GO, KEGG, and Reactome, are available by default for GSEA. Users may customize the database selection. The default adjusted *p-value* threshold for significance is set to 0.05. At this time, GSEA is not supported for metabolomics data, as metabolite-based datasets typically lack sufficient coverage and ranking granularity for reliable enrichment using this method.

### Biological text embedding (Biotext embedding)

To quantify the functional similarity between enriched pathways, MAPA transforms each pathway into a biological text embedding that captures its semantic meaning. This process involves two main steps: pathway information extraction and text-to-embedding conversion.

#### Pathway information extraction

For each enriched pathway, MAPA retrieves relevant textual descriptions from its source database. This biological text serves as the basis for semantic embedding. The extracted information varies slightly depending on the pathway database. (1) Gene Ontology (GO): The pathway text is constructed using the GO term name (title) and its formal definition. (2) KEGG: The pathway description is formed from the pathway title and the narrative summary provided in the KEGG database entry. (3) Reactome: The pathway text includes the pathway name and its general description, typically sourced from the overview or abstract of the Reactome pathway entry. These textual elements are concatenated and preprocessed to form a single, coherent biological text string representing the pathway’s functional context.

#### Text-to-embedding conversion

The constructed biological text is converted into a numerical vector using embedding models accessed via API from OpenAI (text-embedding-3-small). Specifically, MAPA employs an LLM to generate a high-dimensional embedding vector for each pathway. This embedding captures deep semantic information from the pathway’s description, enabling more nuanced comparison than conventional string matching or component overlap. The resulting biotext embeddings serve as a compact and meaningful representation of each pathway’s biological function and are used as input for downstream similarity calculation, network construction, and functional module identification.

### Similarity network construction

Each pathway is first represented as a biotext embedding vector, generated using the OpenAI text embedding model (as described above). These high-dimensional vectors encode the semantic content of pathway names and descriptions, capturing functional meaning beyond simple gene overlap or annotation structure. Pairwise semantic similarity between pathways is then computed using the cosine similarity of their corresponding embedding vectors. Cosine similarity is chosen because it focuses on the angle between high-dimensional vectors over their magnitude^71^. This approach preserves sensitivity to semantic nuances, making it highly effective for assessing functional relationships from pathway descriptions. The result is a pathway similarity matrix, where each entry S_{ij} reflects the semantic similarity between pathway i and pathway j, with values ranging from −1 to 1 (where 1 indicates perfect similarity).

To transform the similarity matrix into a network structure, MAPA applies a similarity threshold (typically user-defined or empirically optimized) to retain only the most meaningful edges. A pathway similarity network is then constructed: (1) Nodes represent individual enriched pathways. (2) Edges represent high-confidence functional similarity between pathway pairs, as determined by cosine similarity exceeding the threshold. (3) Edge weights correspond to the raw similarity scores, which can be optionally retained for weighted network analysis.

This network serves as the foundation for downstream community detection, enabling the identification of functional modules. The thresholding strategy can be tuned to balance network density and biological interpretability.

### Functional module identification

MAPA implements multiple clustering strategies for functional module identification. A functional module is defined as a set of pathways that share related biological functions based on their semantic similarity. MAPA supports three major classes of clustering algorithms: (1) Network-based community detection methods. (2) Distance-based hierarchical clustering methods. (3) Binary-cut, a recursive divisive clustering algorithm designed for similarity matrices^33^. The default method in MAPA is the Louvain algorithm, a graph-based method, but users can choose alternative methods based on their preferences or data characteristics.

#### Network-based community detection

For network-based clustering, MAPA uses the previously constructed pathway similarity network, where nodes represent pathways and edges represent high-confidence similarity relationships based on cosine similarity of biotext embeddings. Users can select from several community detection algorithms provided by the *igraph* R package^56^, such as Louvain (default), Girvan-Newman, Leiden, and Infomap. Effective community detection requires users to select an appropriate similarity threshold to define the network’s edges. The choice of this threshold can be guided by the quality assessment function described below.

#### Distance-based hierarchical clustering

In addition to network methods, MAPA offers a suite of hierarchical clustering algorithms performed using the hclust function that operate directly on the pathway similarity matrix (converted to a distance matrix). Supported linkage strategies include: ward.D, ward.D2, average, complete, single, and others. To generate the final clusters, a similarity threshold should be specified, which is then converted to a height (height = 1 - similarity_threshold) for cutting the dendrogram.

#### Binary-cut clustering

MAPA also supports Binary-cut, a recursive divisive clustering method specifically designed for bioinformatics similarity matrices^33^. This method first generates a dendrogram by recursively splitting the matrix into subgroups at the point of the largest separation in similarity structure. Similar to hierarchical clustering, users must define a similarity threshold to cut the dendrogram, which determines the final set of clusters. The resulting modules can be evaluated using the quality assessment function provided by MAPA.

### Clustering quality assessment

To help users determine the optimal similarity threshold for any chosen clustering method, MAPA provides a dedicated clustering quality assessment function. This function generates a comprehensive report with key statistics to evaluate the appropriateness of the resulting modules for a specific research purpose. The report includes: (1) A summary of clustering results and quality metrics, including the total number of modules, the count and proportion of non-singleton modules, the average silhouette score^72^, the Calinski-Harabasz Index^73^, and the Davies-Bouldin Index^74^. (2) A module size distribution plot to check if modules have reasonable sizes (not too many singleton modules or one dominant large module). (3) A silhouette width plot for visual assessment of cluster cohesion and separation. This output allows users to systematically evaluate different parameters and select the configuration that yields the most statistically robust and biologically coherent functional modules.

### Function module annotation

After functional modules are identified based on pathway similarity, MAPA performs automated and interpretable biological annotation of each functional module using an LLM-assisted pipeline. To inform functional annotation, MAPA gathers relevant biomedical literature that is associated with the pathways in each functional module. This multi-source literature retrieval strategy includes the following steps.

#### Pathway information extraction

For each pathway within a functional module, MAPA retrieves descriptive metadata from the associated pathway databases. For the GO database, the term name and definition are collected. For KEGG, the pathway title and description are collected. For Reactome, the pathway name and general summary are collected. Additionally, pathway components (*e.g.*, genes, proteins, or metabolites) and their linked literature references (*e.g.*, PMIDs) are collected.

#### Literature retrieval from PubMed

MAPA identifies related publications using three complementary strategies: (1) Keyword-based search. Pathway names and molecular components (*e.g.*, gene symbols) are used to query PubMed via API. Reference-based search. Citations associated with each pathway in the databases (*e.g.*, KEGG and Reactome references) are matched to their PubMed entries. (3) User-uploaded literature (optional). Users may also upload a custom set of external publications (PDF format) to supplement automated retrieval, ensuring coverage of unpublished or domain-specific content.

#### Text embedding and similarity ranking

All retrieved publication abstracts (title and abstract) are embedded into high-dimensional vectors using an LLM-based text embedding model (OpenAI). Simultaneously, each functional module is embedded based on the aggregated pathway titles and descriptions it contains. Then, cosine similarity is calculated between the module embedding and all paper embeddings. The top 20 most related publications are selected as candidate literature for annotation.

#### Literature re-ranking and final selection

The top 20 related papers are input into an LLM (*e.g.*, Gemini or GPT-4) for re-ranking and content evaluation. The model selects the top 5 most relevant papers, based on biological coherence and module relevance. These 5 papers (title and abstract) form the final literature set used for downstream summarization.

#### Prompt engineering for functional module summary

The final summarization step uses the selected literature, along with optional study-specific metadata (*e.g.*, phenotypes), to generate structured and interpretable annotations for each module.

Optional experiment metadata input. Users may provide a brief study description or list of associated phenotypes (*e.g.*, aging, diabetes) that are relevant to the context of their omics dataset. This information helps tailor the annotation to specific research questions or conditions.

Prompt-driven summarization. MAPA uses a carefully designed prompt template to instruct the LLM to generate the following structured output:

1. module_name: A concise and biologically meaningful title for the functional module.
2. summary: A paragraph summarizing the key biological processes, pathways, and molecules associated with the module.
3. phenotype_analysis: A section describing how the module may relate to the user-specified phenotype(s), if provided.
4. confidence_score: A numerical score (ranging from 0.00 to 1.00) that reflects how confidently the annotation integrates the pathway content and phenotype context.

Output format. The final output is delivered in structured JSON format, ensuring compatibility with both MAPA’s R-based pipeline and downstream reporting modules. Each annotated functional module includes the pathways, related genes/metabolites, top related publications, and summary text for interpretation and visualization.

### Data visualization

MAPA provides a comprehensive suite of visualization functions to explore and interpret functional module analysis results at multiple biological levels.

#### Pathway enrichment visualization

MAPA creates horizontal bar charts visualizing enrichment results at the pathway, module, and functional module levels. For ORA, users can select x-axis metrics including qscore (-log_10_ adjusted *p-value*), RichFactor, or FoldEnrichment for genes, while metabolite analysis uses qscore exclusively. GSEA results display NES. Bars are colored by database (GO, KEGG, Reactome, HMDB) and scaled by gene/metabolite count, with “straight” (lollipop) or “meteor” (gradient) visualization styles.

#### Similarity network visualization

MAPA visualizes pathway similarity relationships as network graphs at database-specific module or cross-database functional module levels. Node size represents statistical significance (-log_10_ adjusted *p-value* for ORA or absolute NES for GSEA), and edge width indicates similarity strength. Users can filter by module size and display pathway names or LLM-generated module annotations.

#### Multilevel relationship networks

MAPA constructs multipartite graphs showing hierarchical relationships among functional modules, pathways, molecules, and experimental variables. The function supports linear track or circular layouts with customizable node positioning, coloring, and selective inclusion of hierarchical levels.

#### Expression-network integration

MAPA combines relationship networks with expression heatmaps for integrated functional-molecular views. At the pathway level, heatmaps show mean molecular expression with optional word cloud annotations (from pathway descriptions or LLM summaries), where word size represents frequency. At the molecular level, individual expression profiles are displayed with configurable hierarchical clustering.

#### Module-specific analysis

For detailed module exploration, MAPA generates three complementary views: (1) internal similarity networks, (2) statistical bar charts of constituent pathways, and (3) functional word clouds with word size representing cumulative-log10 adjusted *p-values*. This approach enables comprehensive module characterization across network topology, statistical significance, and semantic content.

### Result report

MAPA provides comprehensive output functionality through automated report generation and tabular data export for downstream analysis.

#### Automated report generation

MAPA generates standardized reports in multiple formats (HTML, PDF, Word, Markdown) using R Markdown templates. Reports contain four main sections: (1) analysis parameters and computational settings, (2) enrichment bar plots at pathway, module, and functional module levels showing top 10 results, (3) similarity network visualizations for functional modules across databases, and (4) interpretation tables with LLM-generated summaries when available. The system automatically handles figure placement, table formatting, and conditional content inclusion based on available results with default visualization parameters (p.adjust.cutoff = 0.05, count.cutoff = 5).

#### Data export functionality

MAPA exports all analysis results to individual CSV files for external analysis. Exported data includes enriched pathways from each database (GO, KEGG, Reactome, HMDB), database-specific modules, functional modules, and LLM interpretation results.

### Design of the MAPA shiny application

To make MAPA accessible to a broader range of users, including those without programming expertise, we developed *MAPAShiny*, an interactive web-based interface that encapsulates the full functionality of the MAPA workflow. *MAPAShiny* was built using the Shiny framework in R, a widely used tool for creating dynamic, browser-based applications directly from R code.

The core objective of *MAPAShiny* is to lower the barrier to entry for complex pathway analysis by providing an intuitive, step-by-step user interface. Users can upload their data, select pathway enrichment methods, choose databases (*e.g.*, GO, KEGG, Reactome, SMPDB), set similarity thresholds, and explore functional module annotations, all without writing a single line of code.

Altogether, *MAPAShiny* extends the usability of MAPA by offering a robust, interactive, and transparent environment for pathway-based module analysis, promoting wider adoption in the omics research community.

### Deployment of MAPA and *MAPAShiny*

To support diverse user environments and ensure broad accessibility, we developed and deployed MAPA and *MAPAShiny* in multiple formats. Both tools are implemented as R packages and can be easily installed from GitHub: MAPA (command-line version): https://github.com/jaspershen-lab/mapa. *MAPAShiny* (graphical interface version): https://github.com/jaspershen-lab/mapashiny.

MAPA provides a powerful programmatic workflow designed for advanced users who prefer scripting and batch processing. In contrast, *MAPAShiny* is built as a user-friendly interface, leveraging MAPA’s backend to support interactive, step-by-step analysis via an accessible web interface. This dual design allows MAPA to serve both bioinformaticians and experimental biologists with different levels of computational expertise.

To further streamline installation and ensure cross-platform compatibility, we also provide Dockerized versions of both MAPA and *MAPAShiny*, available at: https://hub.docker.com/r/jaspershenlab/mapa. Using Docker, MAPA can be deployed consistently across different operating systems, including macOS, Windows, and Linux, as well as on local servers, personal computers, and cloud environments. The Docker image encapsulates all necessary dependencies and configurations, enabling reproducible, environment-independent analysis.

In addition to local and Docker-based deployments, we also maintain a fully hosted online instance of *MAPAShiny*, accessible at: https://mapashiny.jaspershenlab.com. This web-based deployment allows users to explore and execute MAPA analyses immediately, without installing any software. Comprehensive tutorials and deployment instructions for all versions are available at: https://www.shen-lab.org/mapa-tutorial.

Together, these deployment options provide a flexible, scalable, and reproducible ecosystem for functional module analysis, empowering researchers to apply MAPA across a wide range of study designs and computational settings.

### General statistics

All data processing, statistical analyses, and visualizations were conducted using R version 4.5.0 (https://www.r-project.org/) within the RStudio environment. A comprehensive list of R packages utilized in this study is provided in the **Supplementary Note**. Statistical significance was assessed using the Wilcoxon rank-sum test implemented via the *ggsignif* package and *wilcox.test* function. Multiple hypothesis testing was corrected using the False Discovery Rate (FDR) method (*p.adjust* function). Effect sizes were calculated to quantify the magnitude of the difference between groups. Specifically, we used the non-parametric statistic Cliff’s delta (δ), which ranges from −1 to 1, where 0 indicates stochastic equality and ±1 indicates complete dominance by one group^75^. Calculations were performed with the cliff.delta function from the *effsize* R package. Dimensionality reduction was performed using Principal Component Analysis (PCA) via the *prcomp* function and Uniform Manifold Approximation and Projection (UMAP) implemented through the *uwot* package. Figure icons were obtained from iconfont.cn and are used under the terms of the MIT License for non-commercial purposes (https://pub.dev/packages/iconfont/license).

### Assessment of pathway redundancy

To evaluate redundancy, we analyzed widely used pathway databases both within-database and across-database. Our within-database analysis focused on the Gene Ontology database. For the across-database comparison, we selected the metabolite databases KEGG, SMPDB, and Reactome.

#### Pathway redundancy within the Gene Ontology database

We assessed the internal redundancy within the human Gene Ontology database. GO annotations were retrieved from the *org.Hs.eg.db* R package (source data stamp: 2025-02-06). Semantic similarity scores between GO terms were calculated separately for the BP, MF, and CC sub-ontologies. This calculation was performed using the Wang method, which considers the topology of the GO graph structure, as implemented in the *GOSemSim* package. The distribution of these similarity scores was then visualized using histograms. To further visualize the redundancy structure of terms with high similarity, we constructed networks based on term pairs with a semantic similarity score greater than 0.9. The networks were built using the *tidygraph* package. Subsequently, we applied the Girvan-Newman community detection algorithm, via the *igraph* package, to cluster the terms into distinct modules. The final networks were visualized using the *ggraph* package.

#### Pathway redundancy across databases

To evaluate redundancy across KEGG, SMPDB, and Reactome metabolite pathways, we first retrieved the original datasets-acquiring human-specific metabolite pathways from KEGG and SMPDB with the *metpath* package^76^ and extracted Reactome human pathways from ChEBI to the all-pathways hierarchy file. After that, we harmonized all metabolite identifiers by mapping the native IDs, like ChEBI IDs, to HMDB IDs via the Tidymass MS1 metabolite database^68^ and removed duplicate HMDB IDs. Finally, we quantified the similarity between pathways by calculating pairwise Jaccard coefficients on the resulting HMDB ID sets.

### Evaluation of the biotext embedding similarity

To rigorously assess the performance of the biotext embedding similarity method developed in MAPA, we conducted systematic benchmarking against multiple established methods commonly used for measuring pathway similarity. Our goal was to determine how effectively biotext embeddings capture functional relationships between pathways compared to conventional strategies based on pathway content or ontology structures.

We benchmarked biotext embedding against several established methods, each representing a distinct paradigm for pathway similarity assessment.

#### Component-Based Similarity

This method quantifies pathway similarity based purely on overlap in molecular components (*e.g.*, genes, proteins, metabolites). Four measures were employed.

1. Jaccard Index. The Jaccard Index is defined as:

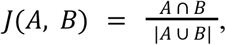

where A and B are the number of molecules annotated to each pathway. This metric ranges from 0 (no shared components) to 1 (identical component sets). It emphasizes proportional overlap but can be biased toward larger pathways.
2. Overlap coefficient. The overlap coefficient is calculated as:

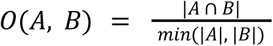 This measure corrects for differences in pathway size by normalizing the intersection over the smaller pathway. It is particularly useful for detecting high similarity between pathways of disparate sizes.

The other two component-based similarity methods are Kappa^77^ and Dice^78^, which can be found in the **Supplementary Note**.

#### Semantic Similarity

Semantic similarity methods capture hierarchical relationships within controlled vocabularies, particularly applicable to GO terms. The Wang method quantifies similarity by considering the graph structure of GO, including the relationships between terms and their ancestors in the ontology. It calculates the semantic contribution of all ancestor terms shared between two GO terms, assigning weights based on proximity in the ontology graph. This method can capture subtle biological relationships not evident through component overlap alone. However, it is limited to GO-based analyses and cannot be applied to pathway databases lacking formal ontology structures (*e.g.*, KEGG or Reactome)^55^. Semantic similarity was calculated using the *GOSemSim* R package, employing the “Wang” method for GO Biological Process terms. Other semantic similarity methods used in this study include Resnik^79^, Rel^80^, Jiang^81^, and Lin^82^, which can be found in the **Supplementary Note**.

#### Evaluation strategy

To assess the performance of different similarity metrics, we applied each approach to the same benchmark dataset, referred to as the curated pathway dataset 1. Because semantic similarity measures are applicable exclusively to GO terms, we first evaluated our biotext embedding method alongside component-based methods by calculating pairwise similarity scores for all pathways in the full dataset. Subsequently, we removed non-GO pathways from the curated pathway dataset 1 and recalculated similarity scores among the remaining GO terms using biotext embeddings, component-based methods, and semantic similarity metrics. This two-step evaluation allowed us to fairly compare methods across both general pathway datasets and GO-specific contexts.

### Evaluation of clustering methods

To evaluate the performance of different clustering algorithms in identifying functional modules, we benchmarked several methods using a curated pathway dataset containing known ground truth module assignments (curated pathway dataset 1). This dataset included both GO terms and pathways from other databases, allowing us to assess clustering performance under diverse biological contexts.

We tested three categories of clustering methods: (1) Graph-based community detection algorithms, including eleven methods (fast greedy, Louvain, walktrap, InfoMap, label propagation, leading eigenvector, Leiden, edge betweenness, optimal, spinglass, and fluid communities) implemented in the *igraph* R package. (2) Distance-based clustering algorithms, including hierarchical clustering with eight linkage strategies (ward.D, ward.D2, single, complete, average, mcquitty, median, centroid) using the *hclust* function, as well as K-means, HDBSCAN, Affinity Propagation, Mean-Shift clustering, and Gaussian Mixture Models, applied to pathway similarity or distance matrices. (3) Binary-Cut, a recursive divisive clustering method optimized for bioinformatics similarity matrices, implemented using the *simplifyEnrichment* R package.

#### Graph-based methods

For graph-based clustering, we constructed similarity networks by applying various cutoff thresholds (ranging from 0.2 to 0.9 in increments of 0.01) to filter edges based on pairwise similarity scores. We evaluated eleven community detection algorithms implemented in the *igraph* R package: fast greedy, Louvain, walktrap, InfoMap, label propagation, leading eigenvector, Leiden, edge betweenness, optimal, spinglass, and fluid communities. Most algorithms operated directly on similarity weights, while the edge betweenness algorithm required distance measures and used converted weights calculated as distance = 1 - similarity. These algorithms automatically determine the optimal number of clusters based on modularity maximization rather than allowing direct specification of cluster numbers. To enable fair comparison across methods with target cluster numbers ranging from 2 to 43, we systematically varied similarity cutoffs and retained only the result with the highest ARI score when multiple cutoffs produced the same number of clusters for a given algorithm.

#### Distance-based methods

We applied multiple distance-based clustering approaches to the pathway similarity matrix. For hierarchical clustering, we converted the similarity matrix to a distance matrix using distance = 1 - similarity and systematically tested eight linkage methods (ward.D, ward.D2, single, complete, average, mcquitty, median, centroid) across all possible cluster numbers from 2 to 43 using the *hclust* function with *cutree* to obtain desired cluster counts. K-means clustering was performed on similarity matrices, directly specifying cluster centers from 2 to 43 with 50 random starts (nstart = 50) for robust optimization. For methods that do not allow direct cluster number specification—including HDBSCAN (with minimum points parameters from 2 to 43), Affinity Propagation (with preference values from 0.2 to 0.9), Mean-Shift clustering (with bandwidth parameters from 0.05 to 1.0), and Gaussian Mixture Models (with component numbers from 2 to 43), we systematically varied their respective parameters and retained only the configuration with the highest ARI score when multiple parameter settings yielded the same number of clusters.

#### Binary-Cut

Binary-Cut clustering was performed using the *simplifyEnrichment* R package with systematic evaluation of cutoff values ranging from 0.5 to 0.95 in increments of 0.01. Since Binary-Cut automatically determines the number of clusters based on the similarity structure rather than allowing direct specification, we applied the *binary_cut* function with try_all_partition_fun = TRUE across all cutoffs and calculated ARI scores against ground truth labels. When multiple cutoffs produced the same number of clusters, we retained only the result with the maximum ARI score to enable fair comparison with other methods across the target range of 2 to 43 clusters.

#### Quantification of consistency between clustering results and ground truth

To quantify agreement between the clustering results and the known ground truth, we computed the Adjusted Rand Index (ARI), a metric that assesses clustering similarity while correcting for chance. Higher ARI values indicate stronger concordance with the curated modules. ARI is calculated using the *adjustedRandIndex* function from the *mclust* R package.

#### Statistical validation of clustering performance

To establish the statistical significance of clustering performance differences, we conducted permutation tests comparing Louvain graph-based clustering and hierarchical clustering (ward.D2 linkage) using the Adjusted Rand Index (ARI) as the evaluation metric. We performed a paired permutation test for dependent ARI scores, where under the null hypothesis of equivalent performance, cluster assignments for any pathway could be interchangeably swapped between methods. We simulated this by randomly swapping cluster labels between Louvain and hierarchical methods for each pathway with probability 0.5 across 9,999 permutations, then calculating ARI differences for each permutation. The *p-value* represented the probability of observing an ARI difference as large or larger than the actual observed difference under the assumption of equivalent performance. All analyses were implemented in R using the adjustedRandIndex function from the *mclust* package, with statistical significance assessed at α = 0.05.

### Negative control experiment with random modules

To validate that MAPA’s annotation system assigns higher confidence to biologically coherent modules rather than arbitrary groupings, we designed a negative control framework based on randomized pathway modules. This approach preserves the structural properties of real modules while removing biological context, enabling a controlled assessment of MAPA’s semantic discrimination.

#### Random module generation

For each curated module containing at least two pathways, we constructed a corresponding random module by sampling the same number of pathways from the pool of all remaining pathways outside the original module. This procedure ensured that random modules were size-matched to real modules but lacked functional coherence.

#### Analysis and comparison

Both real and random modules were processed using the same pipeline. We first computed the average intra-module similarity based on biotext embeddings, and then applied MAPA’s LLM-based annotation to obtain module-level confidence scores. For both metrics, statistical comparisons between real and random modules were conducted using the Wilcoxon rank-sum test.

### Human expert annotations and comparison with MAPA

To evaluate the quality and interpretability of MAPA-generated functional module annotations, we conducted a structured human expert comparison.

#### Expert recruitment and instructions

Five domain experts with backgrounds in molecular biology, biochemistry, and/or computational biology were recruited for this task. Each expert was provided with a list of functional modules, including the pathway names and descriptions for each module, identical to the prompt used for MAPA’s LLM-based annotation (**Supplementary Note**). This ensured that the experts received the same instructions and input information as MAPA. Specifically, the instruction includes pathway information and instructs the experts to generate a biological module name, functional summary, and confidence score based on these inputs. Experts can search databases and literature, but they’re not allowed to use AI tools.

#### Annotation embedding conversion

To enable quantitative comparison of human- and MAPA-generated functional annotations, we converted all annotation texts, both module names and functional summaries, into high-dimensional vector representations (text embeddings). We used OpenAI’s text-embedding-3-small model, which maps each input text into a 1536-dimensional embedding space that captures semantic similarity between texts.

#### MAPA vs. expert comparison

To compare MAPA annotations with expert interpretations, we applied Principal Component Analysis (PCA) to the combined set of text embeddings from both sources for each module. All embeddings were standardized prior to dimensionality reduction to ensure equal contribution of each feature to the overall variance. The top two principal components (PC1 and PC2) were retained for two-dimensional visualization, allowing us to examine the semantic clustering of MAPA-generated outputs versus expert annotations. Modules with missing or incomplete expert annotations (*e.g.*, labeled “Not Available”) were included in the analysis.

### Evaluation of MAPA’s reproducibility

The reproducibility of the MAPA workflow was evaluated over 15 independent runs on the curated pathway dataset 1. Clustering consistency across the runs was quantified using the pairwise Adjusted Rand Index (ARI), calculated using the *adjustedRandIndex* function from the *mclust* R package. To assess annotation stability, the LLM-generated functional summaries were first converted to embeddings using the OpenAI model “text-embedding-3-small”. Finally, we calculated the pairwise cosine similarity between annotations for the same functional module across different runs (to measure consistency) and between different functional modules (to measure specificity).

### Benchmarking MAPA with other existing methods

We compared *MAPA* with three representative established methods: *enrichplot*^36^, *PAVER*^28^, and *aPEAR*^26^. The comparison evaluated two key aspects: functional module identification and annotation ability. Input data requirements, clustering algorithms, and annotation methods for these tools are detailed in **Supplementary Table 1**. We used the curated pathway dataset, comprising 44 pathways from GO, KEGG, and Reactome databases. Since some methods require enrichment results as input, we treated the selected pathways as enriched and generated an artificial enrichment object using the enricher function from the *clusterProfiler* package.

#### Clustering quality comparison

Several methods provide parameters for optimizing clustering results. For *enrichplot*, we tested k-means clustering (default) across all possible cluster numbers (2-43) and selected the configuration with the highest Adjusted Rand Index (ARI). For *aPEAR*, we evaluated all supported clustering algorithms (Markov chain and hierarchical clustering; spectral clustering was excluded due to a known issue). For *PAVER*, hierarchical clustering with dynamicTreeCut was applied, with maxCoreScatter optimized from 0.01 to 1.0 (step size 0.01). Clustering results with optimal ARI were visualized using the *circlize* package.

#### Functional module annotation comparison

To evaluate annotation quality, we applied each method’s annotation functions to the ground truth clustering results. *enrichplot* generates module labels using word frequency analysis of pathway names, with word positioning based on average positions in pathway descriptions. *aPEAR* selects representative pathways using PageRank, while *PAVER* selects the pathway whose embedding has the highest cosine similarity with the average embedding of the module. Both methods generate annotations by combining pathway names and database descriptions. *MAPA* uses LLM-generated functional module names and summaries. All annotations were converted to embeddings using the text-embedding-3-small model provided by OpenAI and compared against expert annotations via cosine similarity.

### Applying MAPA to aging-related multi-omics data

We applied the MAPA framework to analyze functional alterations in aging using publicly available multi-omics datasets^3^. First, we analyzed age-dependent genes identified from bulk RNA-seq data spanning 78 tissues. For each tissue, we compiled a list of dysregulated genes, including both upregulated (cluster U) and downregulated (cluster D). We then performed pathway enrichment analysis on each gene list. Among 78 tissues, 56 tissues with significantly enriched pathways (adjusted p-value < 0.05 and more than five annotated genes per pathway) were retained for further analysis. Using MAPA’s default parameters (Louvain clustering, similarity cutoff = 0.55), we clustered the enriched pathways for each of the 56 tissues, yielding 396 distinct functional modules. To quantify the directional trend of each module with age, we calculated a Dysregulation Index defined as follows:

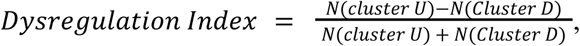

Where *N*(*cluster U*) is the number of upregulated genes and *N*(*cluster D*) is the number of downregulated genes within a given functional module. A higher index indicates that the module is predominantly upregulated during aging, whereas a lower index indicates downregulation.

To identify functionally conserved modules across different tissues, we further clustered the 396 functional modules based on their LLM-generated names and summaries using hierarchical clustering (ward.D2 method, similarity cutoff = 0.55). This process identified 162 meta-modules. For each meta-module, we generated a comprehensive functional annotation by synthesizing the summaries and tissue origins of all its constituent modules using an engineered LLM prompt (**Supplementary Note**). To interpret these results, we visualized the Dysregulation Index for each functional module, organized by meta-module, using a heatmap generated with the *ComplexHeatmap* R package.

Finally, we extended this MAPA workflow to analyze other aging-related datasets, including age-dependent genes from single-nucleus RNA-seq of the liver and brain frontal lobe, as well as age-dependent proteins and metabolites from plasma proteomics and metabolomics data. The resulting functional module networks generated by MAPA are in **Supplementary Figures 9-11**.

## Code availability

All software development, data processing, and statistical analyses were conducted using R version 4.4.1 and its associated packages. The complete source code for MAPA and *MAPAShiny* is freely available at https://github.com/jaspershen-lab/mapa and https://github.com/jaspershen-lab/mapashiny, respectively. All scripts used for data analysis, processing, and visualization in this study can be accessed at https://github.com/jaspershen-lab/mapa_manuscript. Comprehensive tutorials for both MAPA and *MAPAShiny* are provided at https://www.shen-lab.org/mapa-tutorial/, and ongoing updates and maintenance information are available at https://www.shen-lab.org/mapa-website/. The online version of *MAPAshiny* is available at https://mapashiny.jaspershenlab.com.

## Data availability

All the datasets analyzed in this study and used as demo data were obtained from the previously published studies^3,83^. All other source data supporting the findings are provided as **Supplementary Data**.

## Acknowledgments

This work was supported by start-up funding provided to Xiaotao Shen from the Lee Kong Chian School of Medicine and the School of Chemistry, Chemical Engineering, and Biotechnology at Nanyang Technological University, Singapore.

## Author contributions

X.S. conceived the project and provided overall supervision. X.S., Y.G., and F.Z. collaboratively developed the methodology and software. *MAPAShiny* and its Docker implementation were developed by X.S., Y.G., and Y.L. Documentation and tutorials of MAPA were prepared by Y.G., X.S., and Y.S. Case study analyses were conducted by Y.G., C.W., and X.S., who also generated all figures. The manuscript was written by X.S., C.W., Y.G., F.Z., and Y.L., with all authors reviewing and contributing to the final version.

## Additional information

Correspondence and requests for materials should be addressed to X.S.

